# Transient protein phosphorylation promotes disease tolerance to sepsis

**DOI:** 10.64898/2026.06.09.731055

**Authors:** Wolfgang Vivas, Johannes Roth, Katharina Willman, Hjalmar R Bouma, Franziska Röstel, Karen Dlubatz, Gianna Hirth, Nadine Pömpner, Therese Dau, Emilio Cirri, Sheng Zhang, Mirko Peitzsch, Triantafyllos Chavakis, Michael Bauer, Gianni Pannagiotou, Luis F. Moita, Sascha Schäuble, Sebastian Weis

## Abstract

One of the enduring paradoxes of sepsis is that organs fail despite little evidence of irreversible tissue injury. Emerging evidence suggests that this state reflects a regulated metabolic shutdown within host tissues, yet whether such hypometabolism contributes to pathology or promotes survival remains unclear. This phenomenon resembles torpor, a physiological state of profound hypometabolism induced by environmental stress and mediated through reversible protein phosphorylation. Septic hypometabolism is identified here as a conserved tissue-specific metabolic adaptation which is characterized by transient activation of Glycogen Synthase Kinase (GSK)3β. This activation reduced disease severity of bacterial sepsis without affecting the hosts pathogen burden, indicating that GSK3β activity promotes disease tolerance to infection. Consistent with these findings, plasma signatures associated with GSK3β inhibition correlated with worse clinical outcomes in patients with sepsis. Together, these results define septic hypometabolism as a torpor-like response and identify reversible phosphorylation as a key mechanism governing host adaptation to severe bacterial infection.

## INTRODUCTION

Numerous animal species, including reptiles, and certain birds and mammals, have the physiological capacity to reduce their overall metabolic rate and lower energy expenditure. This phenomenon referred to as torpor (short term) and hibernation (long term) allows survival in extreme environments that would otherwise not be compatible with life ^1–3^. It is characterized by a controlled reduction in metabolic rate and core body temperature that is maintained until more favorable external conditions are reached ^4^. Torpid mammals can dramatically decrease their metabolic rate by 95% and reduce vital parameters including the core body temperature down to 5°C ^5^. This is associated with lower mitochondrial respiration and ATP synthesis ^6^. An imbalance, *i.e.*, deficiency between metabolic demands and energy availability can be fatal. Therefore, torpor is subject to stringent regulation, with entry, maintenance and arousal forming tightly coordinated biological processes. Transient protein phosphorylation of protein has been proposed as a key mechanism underlying the reversible physiological state of torpor ^5,7^. This notion is supported by data showing that for example, the activities of enzymes involved in carbohydrate catabolism, including glycogen phosphorylase, hexokinase, and pyruvate dehydrogenase, are reduced during torpor via reversible phosphorylation events ^8,9^. In addition, the kinase activity of central signaling proteins, such as AKT and phosphatidylinositol 3-kinase, are downregulated in hibernating ground squirrels^10^. While the molecular components allowing torpor are conserved between species, mice and humans do not enter a hibernating state under physiological conditions.

Sepsis is a severe and deadly clinical syndrome and defined a decade ago as an infection-associated organ dysfunction ^11^. Increasing amounts of experimental evidence supports the notion that the capacity to appropriately adapt tissue energy metabolism during infection is a key defense mechanism that reduces disease severity and promotes survival ^12–15^. This metabolic adaptation can be afforded in a pathogen-load independent way via promoting a defense mechanism that is referred to as disease tolerance to infection ^12–16^. Notably, the adaptive metabolic responses that occur during severe infections and sepsis closely resemble the physiological changes seen in hibernating animals. Patients with sepsis have reduced oxygen consumption with metabolic rates returning to baseline upon recovery from infection^17^. This indicates that hosts can reduce their metabolic rate in a reversible manner. Torpid animals also show signs of organ and mitochondrial dysfunction ^12,18,19^, which can potentially serve as an adaptive response to preserve energy. Common features between torpor and sepsis-induced hypometabolism are also found at the molecular level. These include, *e.g.*, the activation of the unfolded protein response ^20–24^ and the metabolic stress response via the forkhead box transcription factor (FOXO) ^25,26^. The same genetic programs are proposed to limit pathology and afford tissue damage control in severe infections ^26^.

Yet, whether torpor in sepsis constitutes an adaptation to optimize energy expenditure or this is the core of the maladaptive response resulting in septic organ dysfunction is unknown. In this study we investigated the hypothesis that infected hosts can reversibly reduce their metabolic rate as an adaptive response to improve survival. Our data shows that early metabolic adaptation in sepsis was regulated by transient phosphorylation of proteins, which was associated with a lower disease severity and better outcome. This involved a transient activation of Glycogen Synthase Kinase (GSK)3β as a general feature in the early response to sepsis. Its activation lessened disease severity independent of the host’s pathogen load, indicating that GSK3β was necessary to establish disease tolerance to bacterial infection.

## RESULTS

### Severe bacterial infections trigger a conserved hypometabolic response

To study the infection-induced hypometabolism in a clinically relevant setting, we used a model of polymicrobial sepsis along with application of broad-spectrum antibiotics and fluid resuscitation ^13^. The model had a survival rate of approx. 30%-40% matching the sepsis outcome in patients admitted to an ICU (Figure (Fig.) 1A) ^27^. Across eight days of infection (Fig. S1A), septic mice had a higher clinical severity score (Fig. S1B), a larger reduction in body weight (Fig. S1C), and developed a transient hypoglycemia (Fig. S1D) not observed in sham-operated control mice. Infected animals but not controls went rapidly into a hypometabolic state as shown by a reduction in energy expenditure (EE) (Fig. 1B). Two distinct phases of the host metabolic response were identified and are referred to as *torpor-like,* and *recovery*. The *torpor-like* phase lasted from approximately 6 to 24 hours (h) after infection and was characterized by a rapid drop in EE and accompanied by a reduction in oxygen consumption (VO_2_) (Fig. S1E), CO_2_ production (VCO_2_) (Fig. S1F). Mice became hypothermic (Fig. 1C), had reduced total activity (Fig. 1D; Fig. S1H), respiratory exchange ratio (RER) (Fig. 1E; Fig. S1G), and developed anorexia (Fig.1F) -altogether phenocopying key characteristics of natural torpor ^4,28^. This phase was followed by a transition phase that lasted until day 4 or approximately 96 h after infection. Then animals entered a *recovery* phase from sepsis-induced hypometabolism and reverted their organismal hypometabolism showing normal circadian rhythm. When comparing data from non-surviving *vs.* surviving septic animals, we observed that non-survivors had a deeper level of hypometabolism (Fig. 1G), and the magnitude of EE correlated with disease severity (Fig. S1I) and body temperature (Fig. S1J), suggesting that early metabolic suppression is beneficial only within a bounded range. This phenotype was not specific to polymicrobial peritonitis. When mice were subjected to lung infection with *Klebsiella (K.) pneumoniae*, a main causative pathogen of hospital-acquired pneumonia, the same reaction pattern was observed. *K. pneumoniae* infection caused severe disease and was lethal in 32% of the animals (Fig.1H; Fig. S2A-C). Animals with pneumonia showed a similar rapid hypometabolic response (Fig. 1I-M; Fig. S2D-F) and as seen in septic peritonitis, a more pronounced hypometabolic response correlated with disease severity and mortality (Fig. 1N; Fig S2H,I). We also confirmed that septic hypometabolism is conserved between species by using rats that under physiological conditions do not enter hibernation or daily torpor ^29^. Similar to mice, rats developed septic hypometabolism (Fig. S3A-J). Taken together, our data show that severe bacterial infections trigger a conserved hypometabolic response that shows similar characteristics to the physiology of torpor.

**Figure 1.**
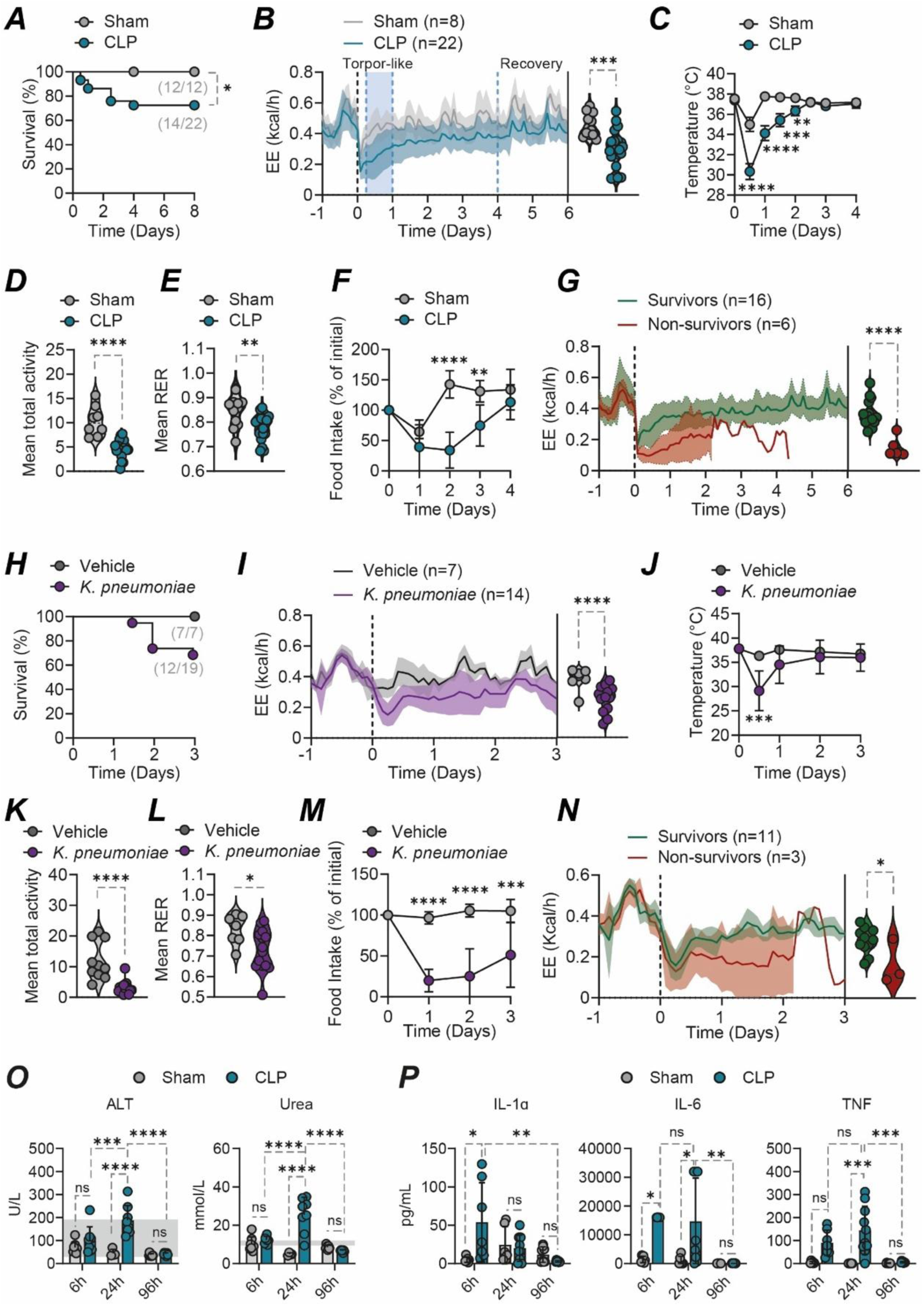
Severe bacterial infection triggers a conserved rapid hypometabolic response. **(A)** C57Bl/6j mice were subjected to either a sham operation or to polymicrobial sepsis via a cecal-ligation and puncture (CLP). Survival of Sham (n=12) or CLP (n=22). **(B)** Energy expenditure (EE) of mice subjected to Sham (n=12) or CLP (n=20). Sepsis induction is depicted in vertical dotted black. Line in curve indicates the mean, and shading denotes ± SD. The graph shows two distinct phases of septic-induced hypometabolism: *torpor-like* and *recovery* (blue dotted line). Violin plot depicts the mean data from individual mice throughout the complete duration of the experiment. **(C)** Body temperature, **(D)** total activity, **(E)**, respiratory exchange ratio (RER), **(F)** and food intake. **(G)** EE of surviving (n=16) *vs.* non-surviving mice (n=6) subjected to CLP. Data pooled from 6 independent experiments. **(H)** Survival of mice subjected to either vehicle (n=7) or pulmonary infection with *K. pneumoniae* (n=19). **(I)** EE, **(J)** body temperature, **(K)** mean total activity, **(L)** mean RER and **(M)** food intake from non-infected or infected mice with *K. pneumoniae*. **(N)** EE of surviving (n=11) *vs.* non-surviving mice (n=3) subjected to *K. pneumoniae*. Data pooled from 4 independent experiments. **(O)** Serology from mice subjected to sham (n=20) or CLP (n=23) during the torpor-like (6 h and 24 h) or recovery (96 h) phase. Grey areas depict reference values from healthy mice. **(P)** Plasma cytokines from mice subjected to sham (n=22) or CLP (n=24) during the torpor-like or recovery phases. Survival in **A** and **H** are represented by Kaplan-Meier plots and the difference between groups was assessed using the log-rank test. Differences in **B**, **D**, **E**, **G**, **I**, **K**, **L** and **N** were assessed with a two-tailed T test. Time course experiments in **C**, **F**, **J**, **M**, **O** and **P** were analyzed by using a two-way ANOVA with post hoc Tukey test. *p < 0.05; **p < 0.01; ***p < 0.001; ****p < 0.0001. <u>Abbreviations:</u> ALT.. alanine aminotransferase; CLP.. cecal ligation and puncture; EE.. Energy expenditure; IL.. interleukin; K.. *Klebsiella*; PCI.. peritoneal contamination and infection; RER.. respiratory exchange ratio (RER); TNF.. Tumor necrosis factor.

We then assessed clinically relevant read-outs in the *torpor-like* and *recovery* phase (Fig. 1B; Fig. S4A). and determined several markers of tissue damage in plasma, including alanine aminotransferase (ALT) and bilirubin for liver damage; urea and creatinine for kidney dysfunction; and LDH as a general damage marker. The extent of organ dysfunction was not different at 6 h between mice subjected to CLP and sham controls (Fig. 1O; Fig. S4B). At this early time point after infection, the ammount of most of these markers were in the range of healthy uninfected mice. Despite no significant differences in LDH plasma levels, ALT, bilirubin, urea and creatinine significantly increased after 24 h in the infected group, (Fig. 1O; Fig. S4B), indicating the progression from infection into sepsis ^30^. Upon recovery, values returned to baseline. Similar to mice subjected to CLP, the levels of most of these parameters was similar at the 6 h time point between the *K.pneumoniae* infected group and controls (Fig. S5A). Since hypometabolism after bacterial infection precedes tissue dysfunction, the data suggest that the onset of septic-hypometabolism is not triggered by organ damage.

We hypothesized that an early systemic cytokine release triggered by the infection could mediate the entry into hypometabolism as suggested earlier ^31,32^. The magnitude of the inflammatory response was assessed by measuring the levels of cytokines in plasma. We found that most of these cytokines were significantly increased at the 6 h time point compared to the sham group (Fig. 1P; Fig. S4C). These inflammatory responses were sustained at the 24 h time point. In the recovery phase, circulatory cytokine levels returned to baseline levels. We also found that the levels of IL-6 and MCP-1 in mice infected with *K.pneumoniae* were significantly increased compared to controls (Fig. S5B).

Notably, elevated plasma IL-6 were found alike in animals subjected to either CLP or *K.pneumoniae* compared to their respective control (Fig. 1P; Fig. S5B). IL-6 is known to modulate the inflammatory response and alter organismal metabolism. Thus, we asked whether IL-6 is the driver that establishes hypometabolism. In a more simplified model, we subjected mice to a non-lethal dose of LPS (Fig. S6A,B,H,I) that triggers a systemic inflammatory response and direct IL-6 release. LPS injection caused several signs of the sickness behavior and torpor in control mice (Fig. S6C-G), as previously reported ^31,33^. When *Il6^-/-^* mice were subjected to LPS, no difference in the hypometabolic response to animals with a physiological expression of *Il6* was observed. Similarly, we observed no significant differences in the LPS-induced hypometabolic response in control mice subjected to either a blocking antibody against IL-6Rα or a control antibody (Fig. S6I-N). Thus, by using two complementary approaches, we conclude that inflammation triggers sickness behavior and a hypometabolic response independently of IL-6.

### Time-resolved and organ-specific metabolic adaptations underlie septic hypometabolism

Our results showed that severe bacterial infection triggers an organismal metabolic response that phenocopies several characteristics observed during torpor. Thus, we hypothesized that distinct changes in cellular metabolism are associated with entry into hypometabolism as observed in torpid animals ^1,34,35^. To test this hypothesis, we performed untargeted and targeted metabolomics in skeletal muscle (rectus femoris), an organ commonly affected during sepsis and torpor ^36–39^, and in the liver as a critical organ controlling organismal metabolic adaptations ^13^. Principal Component (PC) analyses revealed a distinct metabolome in the muscle, with statistically significant differences at the torpor-like phase compared to baseline and the recovery phase (Fig. 2A). We speculated that the distinct metabolome observed during the course of infection would impact the ATP levels in tissues. For this, we performed a flux balance analysis assessing the overall capacity to produce ATP ^40^. We found a higher muscle ATP yield at the septic torpor-like phase (6h) compared to the baseline group and the recovery phase (96h) (Fig. 2B). To understand the underlying mechanisms regulating the predicted ATP yield, we first assessed TCA cycle metabolites with a targeted approach. We found no significant differences in citrate and αketoglutarate, however cis-aconitate level was reduced in the recovery phase compared to baseline and the torpor-like phase (Fig. 2C). Likewise, we found an accumulation of succinate in the muscle in the torpor-like phase that returns to baseline levels upon recovery (Fig. 2C). The accumulation of succinate in the torpor-like phase was accompanied by reduction in the levels of fumarate and malate that prolongs until the recovery phase (Fig. 2C). Indeed, the succinate/malate ratio was elevated during the torpor-like phase and tends to return to baseline levels during the recovery phase (Fig S7B). We observed a reduction in lactate levels during the torpor-like and recovery phases compared to baseline (Fig. S7C). Quantitative pathway enrichment analysis for significantly regulated metabolites revealed a phase-specific enrichment in metabolic pathways. For example, comparing the septic torpor-like phase with uninfected controls, we observed enrichment in stress and redox balance, fatty acid metabolism, and proteostasis and nitrogen handling (Fig. 2D). Conversely, several of these processes were still present at the recovery phase compared to the torpor-like phase, suggesting a transient activation of these pathways during the disease course (Fig. S7D). Indeed, we found a significant reduction of metabolites involved in fatty acid transport into mitochondria, and antioxidant-related compounds during the torpor-like phase compared to baseline animals (Fig. 2E). We also assessed the levels of certain anaplerotic amino acids. While the level of aspartate was elevated during the torpor-like phase, glutamine was reduced compared to the baseline line and the recovery phase groups (Fig. 2E). We found no significant differences in the levels of these metabolites between the baseline and recovery groups. In the liver we also observed a distinct metabolome comparing the septic torpor-like phase with baseline and the recovery phase groups (Fig. 2F). Changes in the liver metabolome in the septic torpor-like phase were associated with a slight, but significant reduction in the ATP yield (Fig. 2G). Similar to skeletal muscle, succinate level was elevated in the liver at the torpor-like and recovery phases compared to baseline levels (Fig. 2H). However, compared to skeletal muscle, the levels of the other evaluated TCA metabolites were reduced in the liver of septic mice during the course of infection compared to the baseline group (Fig. 2H, S7E). Notably, proteomics analyses also revealed a reduced enrichment of the TCA cycle in the liver of *K.pneumoniae* infected mice compared to the uninfected group (Fig. S8A,B). Consequently, the succinate/malate ratio levels were elevated in both the torpor-like and recovery phases (Fig. S7F). We found no differences in lactate levels (Fig. S7G), altogether suggesting a general suppression of the TCA cycle in the liver during the septic torpor-like phase ^41^. Quantitative pathway enrichment analyses revealed enrichment in nitrogen handling, glucose metabolism, and stress and inflammation (Fig. S2I). Similar to skeletal muscle, many of these processes were still enriched in the recovery phase compared to the torpor-like phase (Fig. S7H). We found no differences in the levels of carnitines and aspartate in the liver from mice at the torpor-like phase compared to baseline. However, these were elevated during the recovery phase (Fig. 2J). The levels of antioxidants, glutamate and glutamine were transiently reduced during the course of infection (Fig. 2J). Altogether, similar to what is observed during torpor ^18^, our observations indicate divergent metabolic programs in liver and muscle during the torpor-like and recovery hypometabolic phases. While both tissues show similar patterns during the torpor-like phase into septic hypometabolism, including enrichment in nitrogen handling and antioxidant response, the liver showed a profound reduction in TCA metabolites and anaplerotic amino acids, resulting in a distinct pattern in the ATP yield.

**Figure 2.**
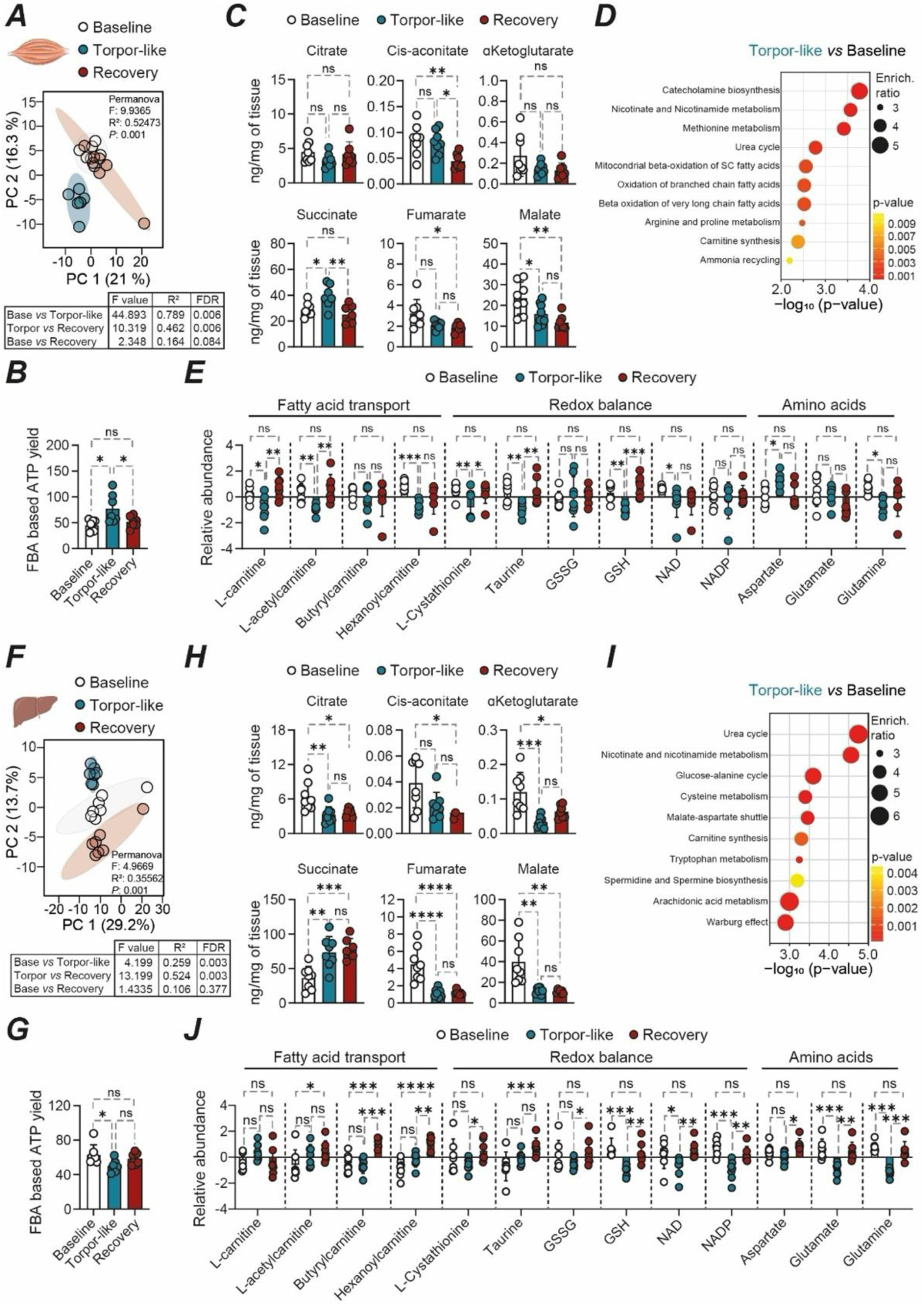
Tissue specific metabolic adaptations underlie septic hypometabolism. **(A)** Principal Component (PC) analysis was generated by unsupervised clustering of LC-MS-based unbiased metabolomics from skeletal muscle obtained from baseline (n=7) and mice in the torpor-like (n=7) and recovery (n=7) phases. The lower panel is shown pair-wise Permanova comparison between groups. **(B)** Flux balance analysis for ATP yield in muscle from baseline (n=7), torpor-like (n=7), and recovery (n=7) phases. **(C)** SMPDB library-based pathway enrichment analysis for significantly regulated metabolites in skeletal muscle from torpor-like phase *vs.* baseline. **(D)** TCA metabolite levels in skeletal muscle from mice at the baseline (n=8), torpor-like (n=8) and recovery phases (n=8). **(E)** Relative abundance of selected metabolites involved in fatty acid transport into the mitochondria, redox balance, and amino acids found in muscle from baseline (n=7), torpor-like (n=7) and recovery (n=7) phases. **(F)** PC analysis was generated by unsupervised clustering of LC-MS-based unbiased metabolomics from liver obtained from baseline (n=7) and mice in the torpor-like (n=7) and recovery (n=7) phases. The lower panel is shown pair-wise Permanova comparison between groups. **(G)** Flux balance analysis for ATP yield in liver from baseline (n=7), torpor-like (n=7), and recovery (n=7) phases. **(H)** SMPDB library-based pathway enrichment analysis for significantly regulated metabolites in liver from torpor-like phase *vs.* baseline. **(I)** TCA metabolite levels in liver from mice at the baseline (n=7), torpor-like (n=7) and recovery phases (n=7). **(H)** Relative abundance of selected metabolites involved in fatty acid transport into the mitochondria, redox balance, and amino acids found in liver at baseline (n=7), torpor-like (n=7) and recovery (n=7) phases. Each point in graphs depicts data from an individual mouse. Data was obtained from 4 independent experiments. Comparison between groups was performed by using one-way ANOVA with post hoc Tukey test. *p < 0.05; **p < 0.01; ***p < 0.001. <u>Abbreviations:</u> ATP.. adenosine triphosphate; CLP.. cecal ligation and puncture; FDR.. false discovery rate; GSH.. glutathione; GSSG.. glutathione disulfide; NAD.. nicotinamide adenine dinucleotide; NADP.. nicotinamide adenine dinucleotide phosphate; ns.. non-significant; PC.. principal component analyses; TCA.. tricarboxylic acid cycle.

### Transient activation of GSK3β is essential to establish disease tolerance to infection

Previous studies indicated that transient phosphorylation of proteins regulate the entry, and arousal from torpor in natural hibernators and in induced torpor ^4,42,43^. Given the phenotypic similarities in septic and natural torpor, we hypothesized that this could be a common regulator. To identify possible kinases that are involved in the initiation of septic hypometabolism, we performed phospho-proteome analysis ^44^ and inferred the most regulated kinases and the corresponding top 3 regulated phosphosites. We performed these studies in liver, skeletal muscle and brown adipose tissue (BAT) ^37,39,45^. At 6 h after infection only four kinases were identified to be activated in the liver of septic animals, namely GSK3β, MAPK8, MTOR, and PRKACA (Fig. 3A). When the same analysis was performed at a later time point, *i.e.* 24 h after infection, the number of affected kinases increased to 22 (Fig. 3B). Hierarchical clustering reveals six major kinase groups (AGC kinases, CAMPK kinases, CMGC kinases, TK kinases, atypical kinases, and others). Notably, GSK3β was the only kinase predicted as activated after 6 h and 24 h. Even more, we found a reduction in the GSK3β activation upon the recovery phase (Fig. S9A). GSK3β is known to regulate the activity of several proteins via phosphorylation, including β-catenin. Phosphorylation of β-catenin targets it for ubiquitination and subsequent proteosomal degradation. As a result, GSK3β activation suppresses the activity of the WNT/β-catenin signaling pathway ^46,47^. Accordingly, we observed downregulation of the “formation of the β-catenin/TCF transactivation complex” in the liver of septic mice at the torpor-like phase (Fig. S9B) which was confirmed by western blot (Fig. S9C). We further aimed at studying the activation of GSK3β using a more simplified model. For this we stimulated precision-cut liver slices from wild type mice with LPS. This resulted in a time-dependent and transient activation of GSK3β, as assessed by the reduction of the inhibitory phosphorylation signal (serine9) in GSK3β (Fig. S9D) and was associated with an increase in the phosphorylation of glycogen synthase, its prototypical downstream target ^48^. This suggests that GSK3β activation is consistent with a liver-intrinsic response that may involve TLR4 signaling.

**Figure 3.**
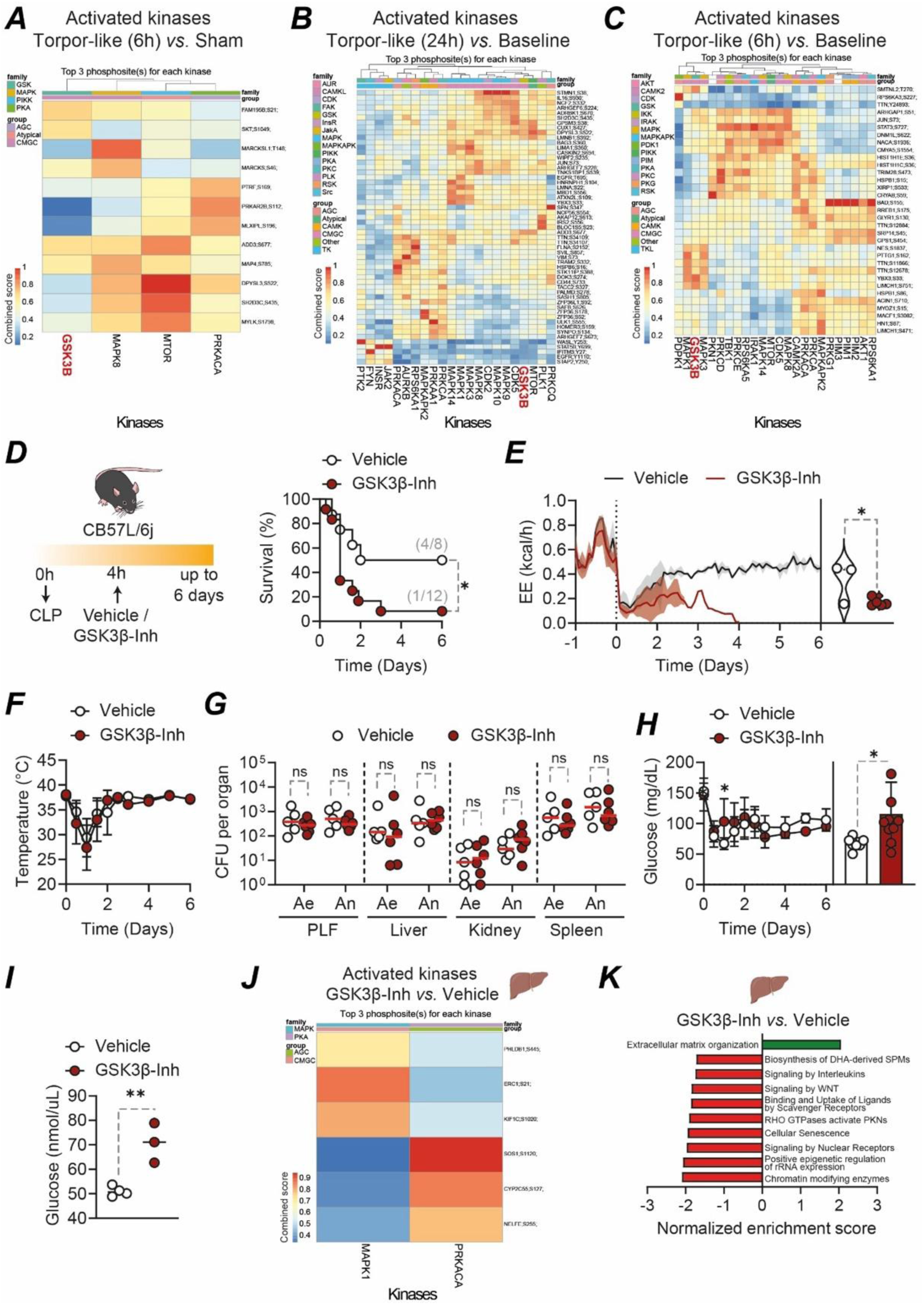
GSK3β activity establishes disease tolerance to infection. Clustered heatmap of the combined kinase-substrate score for the top three phosphosites of all evaluated kinases. The predicted kinases are depicted at the bottom of each heatmap. Analyses were performed from datasets obtained from livers comparing **(A)** CLP (n=8) *vs*. Sham (n=8) mice at the 6 h time-point. **(B)** CLP (n=12) *vs.* Sham (n=8) mice at the 24 h time-point. **(C)** Muscle from septic (n=8) *vs.* control (n=8) mice at the 6 h time-point. **(D)** Survival of mice treated with vehicle (n=8) or the GSK3β inhibitor (SB216763, 25 mg/kg, n=12) 6 h after CLP, data pooled from 3 independent experiments. **(E)** EE from septic mice subjected to vehicle (n=3) or GSK3β inhibitor (n=5). Initiation of surgery is depicted in a vertical dotted black line. The line in the curve indicates the mean, and shading denotes ± SD. Violin plot depicts the mean data from individual mice throughout the complete duration of the experiment. Data pooled from 2 independent experiments **(F)** body temperature from mice subjected to CLP and either vehicle (n=8) or GSK3β inhibitor (n=12), pooled from 3 independent experiments. **(G)** Pathogen loads from septic mice treated with either vehicle (n=5) or GSK3β inhibitor (n=6), pooled from 2 independent experiments. **(H)** Blood glucose levels of septic mice treated with either vehicle (n=8) or GSK3β inhibitor (n=12). The bar plot in the right panel depicts the mean ± SD 24 h after CLP and each point is the value from one mouse, pooled from 3 independent experiments. **(I)** Glucose production from precision-cut liver slices of septic mice subjected to vehicle or GSK3β inhibitor after 6 h of culture, pooled from 2 independent experiment. **(J)** Heatmap from phospho-proteomics analysis in livers depicting predicted activated kinases in CLP mice subjected to GSK3β inhibitor normalized to vehicle-treated mice. **(K)** Gene-set enrichment analysis (GSEA) in livers obtained from CLP mice subjected to vehicle or GSK3β inhibitor. Survival in **D** is represented by Kaplan-Meier plots and the difference between groups was assessed using the log-rank test. Differences between two groups in **E**, **H** and **I** were assessed with an unpaired two-tailed T test. Time course experiment in **F** was analyzed by using a two-way ANOVA. Comparison in **G** was performed with one-way ANOVA with post hoc Tukey test. *p < 0.05; **p < 0.01; ***p < 0.001; ****p < 0.0001. <u>Abbreviations:</u> Ae.. aerobic; an.. anaerobic; CLP.. cecal ligation and puncture; CFU.. Colony Forming Units; EE..Energy expenditure; ns.. Non-significant; PLF.. peritoneal lavage fluid.

Phospho-proteomics in skeletal muscle during the torpor-like phase reveals activation of several groups of kinases including AGC, CAMPK, CMGC and TKL. From these, again GSK3β was activated in septic mice (Fig. 3C). Likewise, we found activation of GSK3β in BAT from septic mice compared to control mice (Fig. S10A). Altogether, our data showed a complex dynamic in the kinome during sepsis, with organ-specific activation patterns that include the common activation of GSK3β during the torpor-like phase.

GSK3β is a ubiquitous central kinase with regulatory functions in glucose metabolism, insulin activity, and energy homeostasis ^49–51^. Since we observed a generalized activation of GSK3β across several organs during the septic torpor-like phase, we treated mice with vehicle or a GSK3β inhibitor (SB216763, 25 mg/Kg ^52^ either 2 h prior CLP or 6 h after CLP (Fig. 3D; Fig. S10B). We used this approach since ubiquitous constitutive deletion of *Gsk3b* is postnatally lethal ^53^. Inhibition of GSK3β increased mortality during polymicrobial sepsis (Fig. 3D; Fig.S10C), suggesting a protective role of GSK3β at the onset of sepsis. Septic mice treated with the GSK3β inhibitor showed a significant reduction in EE (Fig. 3E), and VO_2_ (Fig. S10D) but not in other parameters of host metabolism, including body temperature (Fig. 3F, S10E-J). SB216763-treated mice did not exhibit significant difference in serological markers for liver injury (ALT, bilirubin), kidney dysfunction (urea, creatinine), or general damage (LDH) compared to vehicle-treated mice (Fig. S11A,B). The protection afforded by GSK3β activity was not associated with changes in systemic cytokine release (Fig. S11C) or immune cell composition (Fig. S11D). Likewise, pathogen loads of animals treated or not with the GSK3β inhibitor were indistinguishable, as assessed for aerobic and anaerobic bacteria in different organs 14 h after CLP (Fig. 3G). This indicates that early activation of GSK3β is protective and is required to establish disease tolerance to sepsis.

To test whether GSK3β activity in hepatocytes controls disease severity and organismal metabolism in sepsis, we generated targeted *Gsk3b* knockout in hepatocytes by crossing *C57BL/6 Gsk3b^flox/flox^* mice with *C57BL/6Alb^Cre^*mice. We confirmed a significant reduction of GSK3β in the liver (Fig. S12A). The remaining expression of GSK3β in the *Gsk3b^AlbΔ/Δ^* mice likely stems from non-parenchymal cell origin ^54,55^. When subjected to CLP, *Gsk3b^AlbΔ/Δ^* mice did not develop any phenotype in comparison to control animals with physiological expression of *Gsk3b* (Fig. S12B-F). Likewise, we found no differences in metabolic parameters (Fig. S12G-J). These data suggest that either *i.)* GSK3β signaling in hepatocytes does not have an impact on the disease severity and does not play a role in modulating organismal metabolism during polymicrobial sepsis and/or *ii.)* compensatory mechanisms from other tissues or cells are activated in response to the lack of GSK3β signaling in hepatocytes.

Alterations of glucose metabolism were previously linked to disease severity and mortality during sepsis ^13,14^. In line with this data, we found that septic mice subjected to the GSK3β inhibitor had elevated levels of blood glucose during the early stage of infection compared to vehicle treated mice (Fig. 3H). Elevated glucose levels can be caused by impaired clearance through insulin resistance and/or excess glucose production from the liver via gluconeogenesis or glycogenolysis ^56^. We collected liver from septic mice subjected to either vehicle or SB216763 and performed precision-cut liver slices to quantify glucose output in response to gluconeogenic precursors. Glucose production was higher in liver slices from the GSK3β inhibitor-treated mice with sepsis compared to vehicle-treated septic mice (Fig. 3I). This indicates that glucose output by the liver contributes to hyperglycemia in the GSK3 inhibitor-treated septic mice. However, this was not associated with changes in gene expression of rate-limiting genes for gluconeogenesis or glycogenolysis (Fig. S11E) or with altered glycogen availability (Fig. S11F). Phospho-proteome analysis revealed upregulation of two kinases upon GSK3β inhibition, MAPK1 and PRKACA. The latter is a downstream kinase in glucagon and epinephrine signaling (Fig. 3J) ^56^. Whether this is involved in the observed higher glucose output remains to be established. Gene-set enrichment analyses also revealed suppression of several pathways including in stress responses and metabolic adaptations, including specialized pro-resolving mediators (SPMs) biosynthesis, interleukin signaling, WNT signaling, nuclear receptors, scavenger receptors and regulation of translational capacity (Fig. 3K). Together, this set of data indicate that early activation of GSK3β is a component of the adaptive metabolic response to sepsis. Its inhibition compromises disease tolerance to infection by disrupting systemic energy homeostasis and hepatic glucose output.

### Hypometabolic-associated plasma proteomics correlates to human sepsis severity

We then asked whether the sepsis torpor-like state is associated with a distinct proteomic trait in the plasma ^57^. If so, this would enable the identification of clinically relevant circulating biomarkers which accurately reflect the hypometabolic phenotype, thereby facilitating translation into human medicine. First, we performed plasma proteomics from the mice infected with *K.pneumoniae* and from uninfected control animals. Differential expression analysis identified 82 infection-associated proteins. Of those, 54 proteins were upregulated (FDR < 0.05, Fold Change (FC) > 1.5) and 28 were downregulated (FDR < 0.05, FC < 1.5) (Fig S13A). Gene-set enrichment analysis revealed enrichment only in the downregulated proteins in terms of “Signaling by GPCR”, “Lipoprotein assembly, remodeling, and clearance”, “telomere maintenance”, and “DNA damage/stress induced senescence” (Fig. S13B). We then compared this mouse dataset with three recently published datasets obtained from patients with sepsis ^58–60^ (Fig 4A). We found that 28 proteins in the plasma from *K.pneumoniae* infected mice were also identified in human sepsis datasets (Fig 4B,C,D). Among these, 15 proteins were upregulated (Fig. 4B,D) and five proteins were downregulated (Fig. 4C,D). Further eight proteins showed species dependent abundance (Fig. 4D), indicating species-specific responses during acute severe infections. The overlapping differentially expressed proteins were mapped to protein-protein interaction networks. This confirmed significant interacting modules of acute phase and immune responses, and glyceraldehyde-3 phosphate metabolic process for upregulated proteins (Fig. S13C) and lipoprotein metabolism for downregulated proteins (Fig. S13D).

**Figure 4.**
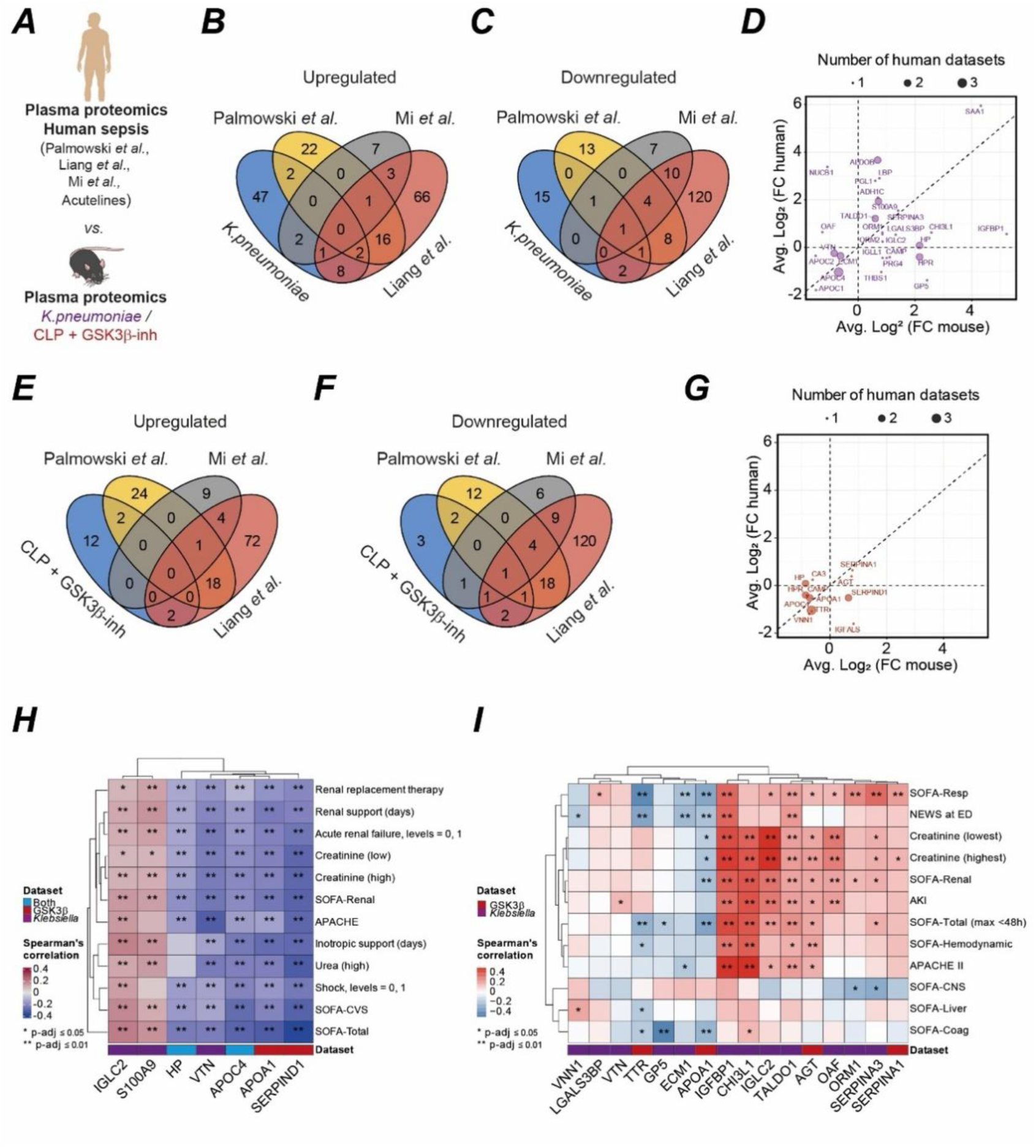
Hypometabolic-associated plasma proteomics correlates to human sepsis severity. **(A)** Plasma proteomics dataset from *K.pneumoniae*-infected mice and GSK3β inhibitor CLP mice were compared to published available datasets of human sepsis ^58–60^. Venn diagram depicting the number and overlaps of differentially **(B)** upregulated and **(C)** downregulated proteins in the *K.pneumoniae* infected mice dataset compared to human sepsis datasets. **(D)** Correlation of differentially expressed proteins from human datasets compared to the *K.pneumoniae* infected mice dataset. Venn diagram depicting the number and overlaps of differentially **(E)** upregulated and **(F)** downregulated proteins in the CLP mice treated with GSK3β inhibitor dataset compared to human sepsis datasets. **(G)** Correlation of differentially expressed proteins from human datasets compared to the CLP GSK3β inhibitor mice dataset. **(H)** Heatmap of correlations between proteins from our mic datasets with clinical variables associated with severe illness in humans and proteins found to be shared in the *Mi et al.* 2024 dataset. **(I)** Heatmap of correlation of significantly expressed proteins in mice datasets with early human sepsis from the Acutelines data- and biobank ^61^. <u>Abbreviations:</u> FC.. Fold change; K.. *Klebsiella*.

Our previous experiments showed that transient activation of GSK3β was essential to establish hypometabolism to polymicrobial sepsis. We then performed plasma proteomics from mice that were subjected to CLP and treated with either GSK3β inhibitor or vehicle in order to understand GSK3β-specific regulation of plasma proteome in the torpor-like phase of sepsis. Differential expression analysis identified 43 infection-dependent proteins with 21 proteins upregulated (FDR < 0.05, log_2_FC> 1.5) and 22 downregulated (FDR < 0.05, log_2_FC< 1.5) (Fig S13E). Gene-set enrichment analysis revealed positive enrichment in “digestion” and negative enrichment terms in “plasma lipoprotein assembly”, “metabolism of lipids”, “cellular senescence”, and “mitotic prophase” (Fig. S13F). We found that 12 proteins in the plasma from CLP mice subjected to GSK3β inhibitor were also identified in human sepsis datasets (Fig 4E-G). Among these, two proteins were upregulated (Fig. 4E, G) and six proteins were downregulated (Fig. 4F,G). Two proteins showed to be specific in human sepsis while two proteins were specific to CLP mice subjected to GSK3β inhibitor (Fig. 4G).

We then asked whether the expression of these proteins in our mouse datasets correlates with the severity of human sepsis and mapped the significantly enriched proteins to clinical human disease severity parameters using the Mi *et al.* dataset ^59^. We found that two upregulated proteins (IGCL2 and S100A9) in the *K.pneumoniae* infected mice were associated with a worse clinical outcome in human sepsis while three proteins (HP, APOC4, and VTN) were associated with better clinical outcome during sepsis (Fig. 4H). Likewise, we found that four of the downregulated proteins in the GSK3β-inhibited mice with sepsis were associated with better clinical parameters in human sepsis (Fig. 4K). Notably, HP and APOC4 were shared between both mouse models. We further analyzed our dataset with dataset of the Acutelines data- and biobank which include samples obtained at the earliest possible time point, *i.e.* the presentation at the emergency room ^61^. Patients’ characteristics of this cohort are provided in *Table 1*. Comparing our *K.pneumoniae* dataset, we found that five proteins (VNN1, LGALS3BP, VTN, GP5, and ECM1) and seven proteins (IGFBP1, CHI3L1, IGLC2, TALDO1, OAF, ORM1, and SERPINA3) were associated with either a better or worse clinical outcome, respectively (Fig. 4I). Notably, two downregulated proteins (TTR and APOA1) in GSK3β-inhibited mice were associated with better clinical parameters. In contrast, two upregulated proteins (AGT and SERPINA1) in GSK3β-inhibited mice were associated to worse clinical severity in human sepsis. This approach allowed us to identify a conserved plasma proteomics signature that is associated with the torpor-like septic phenotype in mice and with disease severity in patients with sepsis. We also found that inhibition of specific regulated proteins by GSK3β lead to a distinct expression pattern of proteins that correlated with a worse clinical severity in septic patients. This provides evidence for torpor-like septic states in humans in relation to infection severity.

## DISCUSSION

Torpor is a regulated physiological response of profound hypometabolism in response to stressful environmental conditions promoting survival ^1,3,4^. Severe infectious diseases are associated with internal stress in which the limited energetic resources need to be managed properly. Infected hosts are forced to adapt their metabolism in a way to balance clearance of pathogens and maintenance of vital organ functions ^26,62^. This includes adaptation of regulated variables such as body temperature or food intake and allows shifting homeostatic parameters to free capacities to fight pathogens while preserving organ function ^63^ a mechanism that we refer to as tissue damage control ^26^. Previous data from animal and clinical studies suggest that a regulated cellular metabolic shutdown could be an underlying conserved response to severe infections ^12,17,31^. However, when maladaptive, *e.g.* when the adaptive response becomes too pronounced or is not reversed appropriately this could result in energetic and metabolic failure of the host, *i.e.* typical traits of sepsis, which correlates with disease mortality ^64,65^. In contrast, an appropriate metabolic adaptation during infection maintains organismal homeostasis of the infected host and underlies disease tolerance to infection ^13,66^. This is in line with our data indicating that the hypometabolic response can be both adaptive and maladaptive since the depth and irreversibility of hypometabolism correlated with disease severity and mortality. It suggests that there is a threshold of hypometabolism severity that turns an initial adaptive response into a mal-adaptive response that becomes detrimental.

Notably, changes in metabolism during infection-associated hypometabolism parallel those observed during torpor ^4,28^ which has been shown to be controlled by transient protein phosphorylation ^42,43^. The data presented in this manuscript suggests that early transient phosphorylation is a critical component of the host response to achieve a hypometabolic state at sepsis onset. We found that a transient activation of glycogen synthase kinase 3β (GSK3β) was a general feature of the early septic response. Originally discovered as an inhibitor of glycogen synthase, GSK3β regulates hepatic glycogen storage and glucose release, thereby determining systemic fuel distribution during stress ^67^. GSK3β activation during early sepsis was an adaptive trait since its inhibition was associated with a deeper septic torpor and resulted in higher disease severity. GSK3β activation acts independently of the cytokine release, and more intriguingly, the hosts pathogen load, hence conferring tissue damage control ^26^. This supports earlier data that highlighted the centrality of glucose metabolism for disease tolerance to sepsis ^13,14^. However, our data contrasts an earlier study showing an inverse infection survival by GSK3β inhibition ^68^. There, the authors did not measure pathogen loads, systemic cytokine release or metabolism rendering the direct assessment with only one comparator difficult. Whether the application of antibiotics in our model was the relevant factor remains speculative. In principle, the inhibition of GSK3β should limit glucose output of the liver. However, our data reveals that during sepsis, a more complex pathophysiological response occurs in which inhibition leads to increased glycemia. Potentially, this could be caused by enhanced gluconeogenesis via the Cori cycle or other compensatory mechanisms, may they be liver intrinsic or not. The ability of GSK3β to control septic torpor extends its known functions beyond cell-injury modulation to encompass systemic energy coordination and survival strategies during infection. Potentially, GSK3β activity links energy, *i.e.* glucose sensing in the liver with systemic metabolic adaptation ^67^. This interesting phenomenon is currently under further investigation.

This study raises several questions. It is not clear if our observations on transient phosphorylation controlling hypometabolism will also apply to infections caused by other classes of pathogens or whether this is restricted to bacteria. Previous data with plasmodium infection, the causative agent of malaria ^69^ or the application of the viral pathogen-associated molecular pattern Poly(I:C) ^31^ however show a comparable energetic response pattern in phenotyping cages which suggests as genera response. Furthermore, it remains to be revealed what is the first signal or combination of signals that is triggered after infection and before the onset of sepsis which results in the transient protein phosphorylation associated with hypometabolism. While our findings suggest translational relevance, the confirmation of a critical role of GSK3β in human sepsis will require dedicated clinical validation. This is hampered by the inability to reach parenchymal organs in severe infections ^70^. We here identified plasma protein surrogate parameters that are conserved in septic patients and mice. These could indicate the early hypometabolic phase of infection and allow for the study of the entry into septic torpor-like response. Previous work had identified a specific neuronal population in the preoptic area of the hypothalamus to control physiological torpor ^71–73^. Future work could delineate cell-type-specific kinase cascades and define how neuroimmune circuits coordinate these pathways. Elucidating these interactions will clarify whether controlled augmentation of septic torpor can be leveraged therapeutically. Notably, induced torpor can slow aging in mice ^74^ which supports the notion of a disease sparing capacity. Ultimately, we did not assess how transient phosphorylation regulates the brain area that controls septic hibernation, since this was not the scope of our work.

In summary, disease tolerance during early sepsis depends on rapid, phosphorylation-driven metabolic adaptation resembling natural torpor. GSK3β serves as one key integrator establishing a mechanistic link between energy sensing and systemic hypometabolism in the septic host.

## Lead contact

Further information and requests for resources and reagents should be directed to and will be fulfilled by the lead contact, Sebastian Weis, MD.

## Materials availability

This study did not generate new unique reagents.

## Data and code availability

- Patients’ characteristics from the Acutelines dataset are provided in *Table 1*.
- Any additional information required to reanalyze the data reported in this paper is available from the lead contact upon request.

## ACKNOWLEDGMENTS

**SW** was supported by the Center for Sepsis Control and Care (CSCC) at the Jena University Hospital. The CSCC was funded by the German Ministry of Education and Research (BMBF No. 01EO1502). **SW** is currently funded by the Deutsche Forschungsgemeinschaft (DFG, German Research Foundation) under Germany’s Excellence Strategy – EXC 2051 – Project-ID 390713860 and by the Deutsche Forschungsgemeinschaft, DFG, project number WE 4971/6-1. **WV** was supported by the Interdisziplinäres Zentrum für Klinische Forschung (IZKF) – Project MSP19. **LFM** is supported by La Caixa Foundation grant LCF/PR/HR23/52430007 and **KW** by an FCT (Portuguese Foundation for Science and Technology) CEEC individual contract (CEECIND/02661/2018). **SaS** and **GP** were supported by the DFG CRC/Transregio 124 “Pathogenic fungi and their human host: Networks of interaction” (DFG project number 210879364, subproject INF). **TC** is supported by the DFG CRC/Transregio 412, subproject B03. The Acutelines data- and biobank infrastructure at UMCG has been established with funds from the University Medical Center Groningen. The authors would like to thank Acutelines and all its participants. The establishment of Acutelines has been made possible by funds from the University Medical Center Groningen. The authors thank Dr. Elisa Jentho and Dr. Dania Martínez Alarcón for critical assessment and discussion of the manuscript. We thank Dr. Karina Xavier and Dr. Vitor Cabral for providing the *Klebsiella pneumoniae* strain MH258.

## DECLARATION OF GENERATIVE AI AND AI-ASSISTED TECHNOLOGIES IN THE WRITING PROCESS

During the preparation of this manuscript, *ChatGPT* (GPT-5.2, https://chatgpt.com) and *Perplexity* (https://www.perplexity.ai/) were used to improve grammar, writing style, and clarity of the text, and to assist with literature searching. All outputs were reviewed and edited by the authors, who take full responsibility for the content of the manuscript.

## AUTHOR CONTRIBUTIONS

Conceptualization, SW; Data Curation, HRB, EC, SZ, MP, SS; Formal Analysis, WV, HRB, EC, SZ, P, SS; Funding Acquisition, WV, LM, SW; Methodology, WV, EC, MP, SS, SW; Project Administration, WV, GH, SW; Resources, all authors; Software, SS; Supervision, GP, MB, LFM, SW; Visualization, WV, HRB, EC, SS; Writing – original draft, WV, SW; Writing – review & editing, all authors.

## DECLARATION OF INTERESTS

The authors declare no conflict of interest.

## STAR METHODS

### KEY RESOURCES TABLE

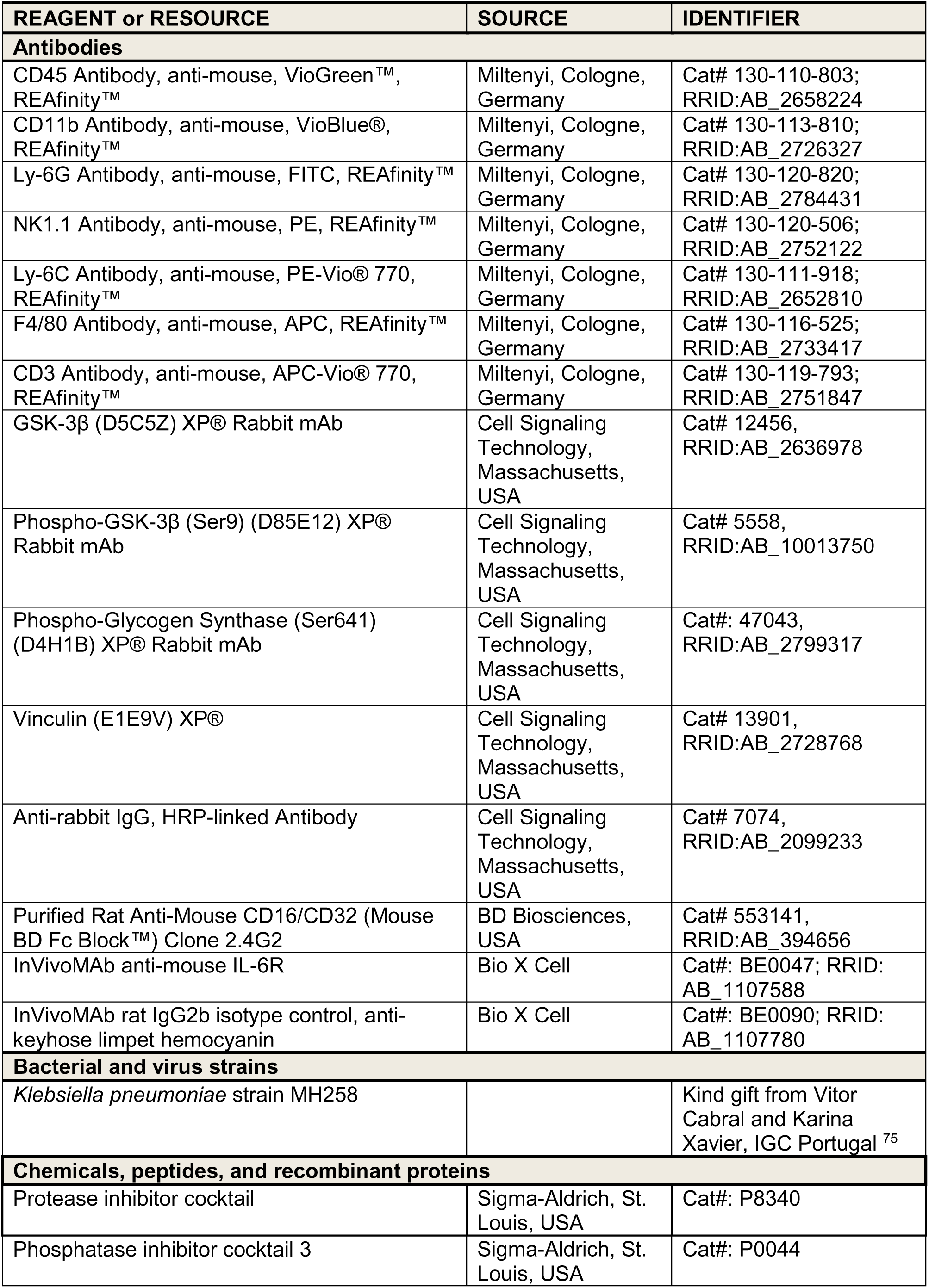
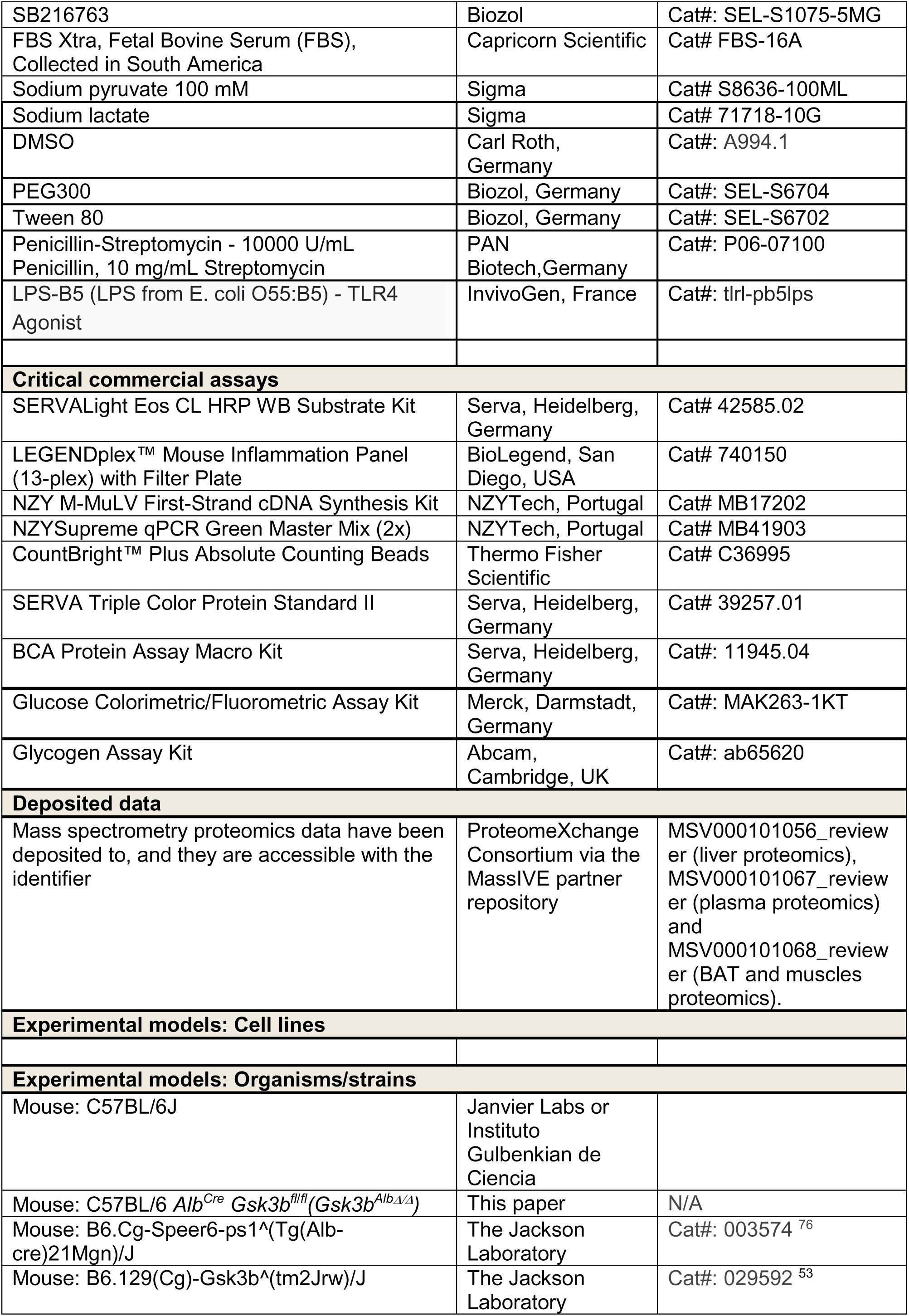
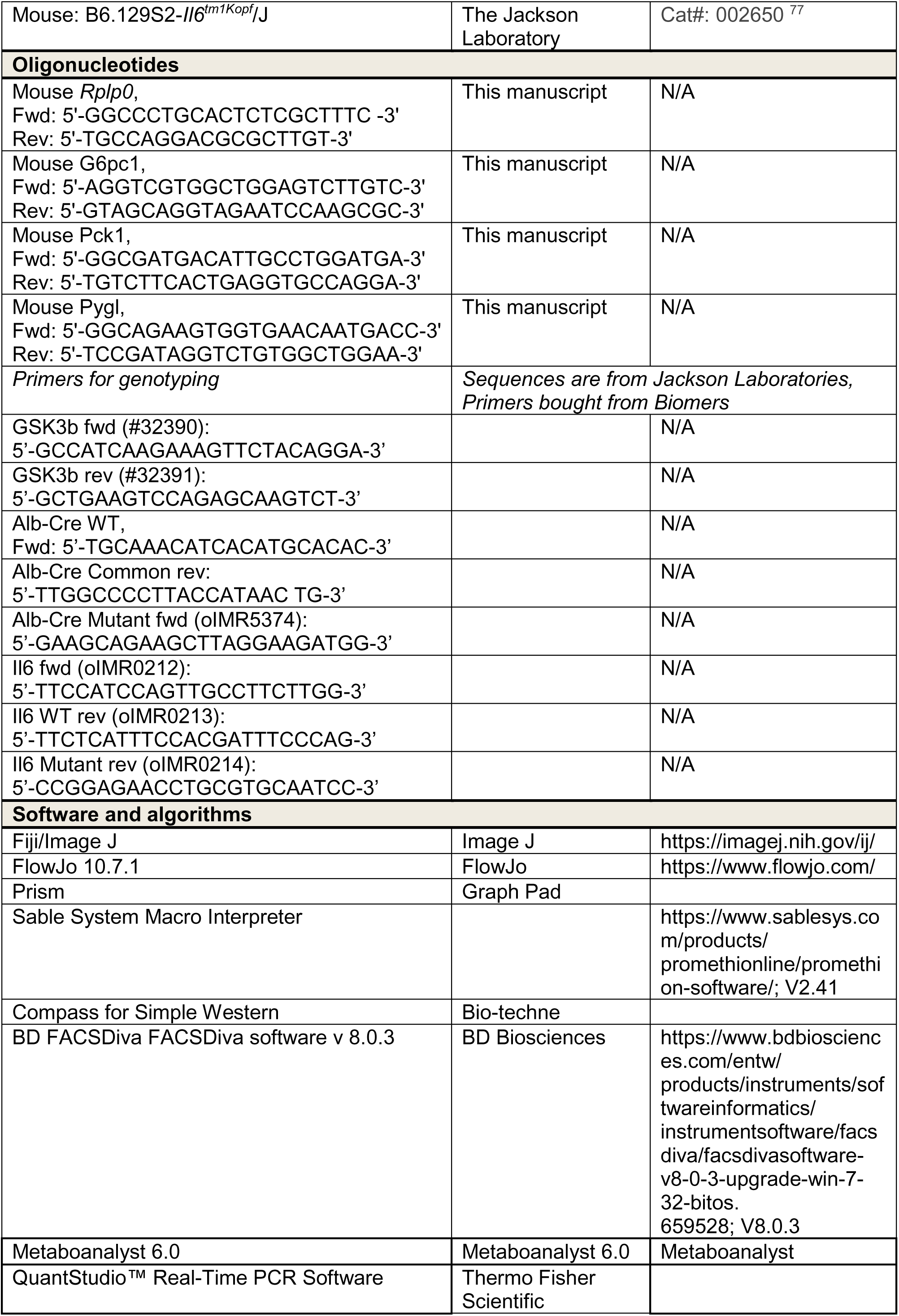
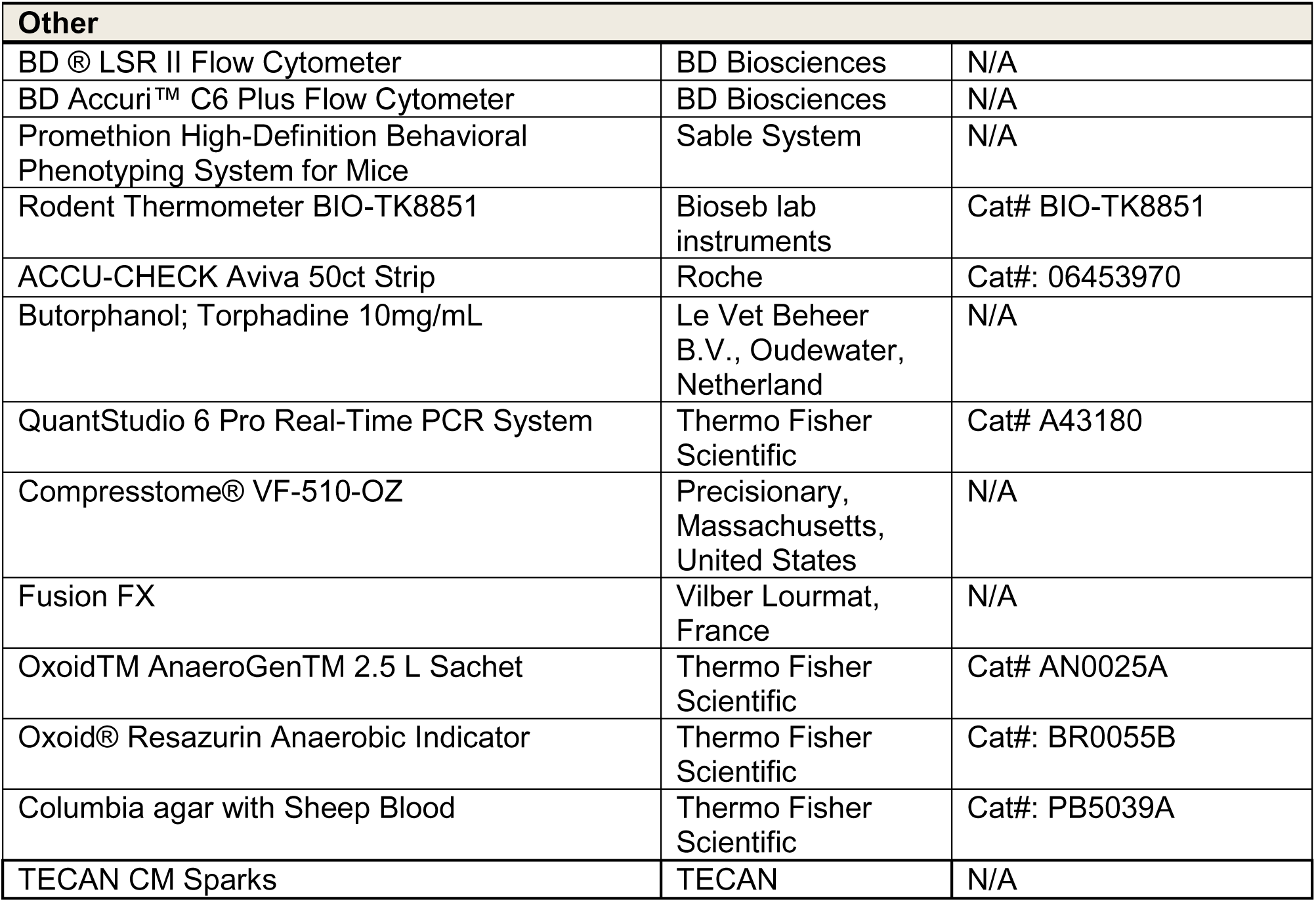

### EXPERIMENTAL MODEL AND SUBJECT DETAILS

#### Animal Experiments

##### Animals

Experimental procedures were ethically reviewed and approved by the Ethics Committee of the regional animal welfare committee (Registration number: UKJ-19-019, HKI-22-004, HKI-22-009, HKI-23-005, HKI-24-001, HKI-25-004, Thuringian State Office for Consumer Protection and Food Safety) or the Instituto Gulbenkian de Ciência (license reference: A011/2019, by Direção Geral de Alimentação e Veterinária. All experiments conducted on animals followed the German legislation on protection of animals, the Portuguese (Decreto-Lei n° 113/2013) and the European (Directive 2010/63/EU) legislations, concerning housing, husbandry and animal welfare. Mice were kept in two different animal facilities, both specific pathogen free (SPF). Animals were group housed (2-5 animals per cage) under controlled temperature (22-24°C) and humidity (40-50%) conditions on a 12 h light/12 h dark cycle. Mice were single-housed only when required by the experiment, *i.e.* indirect calorimetry. Mice (C57BL/6j) and rats (WISTAR RjHan:WI) were fed *ad libitum* with a standard rodent chow diet, containing 57% carbohydrates, 34% protein and 9% fat (Ssniff® Spezialdiäten GmbH, V1554) and had *ad libitum* access to water. Animal groups were formed to have similar numbers of females/males of the same age (12-18 weeks). *Gsk3b^AlbΔ/Δ^*, mice were generated by intercrossing *C57BL/6 Gsk3b^flox/flox^* mice (Jackson Laboratories; stock 029592) with *C57BL/6Alb^Cre^* (Jackson Laboratories; stock 003574). Deletion of *Gsk3b* gene was confirmed in the liver by western blot (Fig. S8A). *Il6^-/-^* mice (Jackson Laboratories; stock 002650) were purchased from the Jackson Laboratories and used directly in experiments after an adaptation period of at least 4 days.

##### Cecal Ligation and Puncture

CLP was performed as previously described ^13^. All experiments were routinely performed at similar times starting at 10-11 AM. Rats were anesthetized with 75 mg/kg / 15 mg/kg Ketamin/Xylazin (*i.p.*). Mice were anesthetized with either 75 mg/kg / 15 mg/kg Ketamin/Xylazin (*i.p.*) or with inhalation of isoflurane (2-5 Vol% for initiation, 1,5-3 Vol% for maintaining narcosis) and periprocedural analgesia with 1 mg/kg butorphanol (*s.c*., given 30-60 min before the operation and 4 h after). Under deep narcosis the cecum was exteriorized, ligated to either 30% or 50% (in all inhibitor and GSK3β-KO experiments) followed by a double puncture with a 23 Gauge needle. After puncture, a small amount of feces was extruded and the cecum was placed back into the abdominal cavity. Sham animals were subjected to the same procedure but without ligation and puncture. Animals were kept on warming mats during operation and narcosis. Eye ointment with dexpanthenol was applied to prevent corneal damage. All animals received 0.9% saline (40 mL/kg; s.c.) once and Imipenem/Cilastatin (25 mg/kg; s.c.) starting 6 h after CLP and every 12 h for a total of three days. Weight, blood glucose levels (ACCU-CHEK Aviva, Roche), and core temperature (Rodent Thermometer BIO-TK8851, Bioseb, France) were assessed. Disease severity was monitored using an established Clinical Severity Score for the entire duration of the experiment. Mice were euthanized when they reached defined end points, i.a. a core temperature 2x below 28 °C or 1x below 25 °C.

##### Klebsiella pneumoniae infection

Pulmonary infection was carried out by using the *Klebsiella pneumoniae* clinical isolate MH258^75^. *K. pneumoniae* was grown in Lysogeny Broth (LB)-media to early exponential stage, harvested, washed and diluted with PBS to perform intranasal application of 1 x 10^8^ colony forming units (CFU) in previously anesthetized mice with isoflurane. Survival was assessed for a total of 4 days. Weight, blood glucose levels, temperature, and disease severity were monitored for the duration of the experiment similar to the CLP experiments.

##### Pharmacologic Approaches

The GSK3β inhibitor SB216763 (Biozol, Ref: SEL-S1075-5MG) was suspended in the vehicle (20%DMSO+40%PEG300+5%Tween80) at a concentration of 2.5 mg/mL. Inhibitor was administered once via an intraperitoneal injection (25 mg/Kg body weight) either 2 h prior or 6 h after CLP. Data obtained from the different time points are described in the figure legends. The vehicle was administered at the same dosage to control mice.

For LPS endotoxemia, mice were injected once intraperitoneally with ultrapure LPS from *Escherichia coli* (InvivoGen, Ref: tlrl-pb5lps) diluted in endotoxin-free PBS (Merck, Ref: TMS-012) at a dose of 5 mg/kg body weight. For experiments with antibody blockage, anti-IL6Ra or the isotype antibody (InvivoGen; Ref: BE0047 and BE0090, respectively) diluted in endotoxin-free PBS was intraperitoneally injected one day before and 4 h after LPS injection at a dose of 10 mg/kg body weight. Vital parameters were measured as described above.

### METHODS DETAILS

#### Serology

Mice were euthanized with ketamine/xylazine overdose or CO_2_ inhalation. Blood was collected via cardiac puncture using a heparinized syringe. Blood samples were stored for 30 min at 4°C and subsequently centrifuged at 2000 g, 10 min, 4°C. The plasma was then collected into a fresh 1.5 mL Eppendorf tube and stored at −80°C until further analysis. Measurements of tissue damage serological parameters were performed by SYNLAB.vet GmbH (Leipzig).

#### Cytokine Measurement

Released cytokines were determined in plasma using the LEGENDplex™ Mouse Inflammation Panel 13-plex (BioLegend, San Diego), according to manufacturer instructions.

#### Promethion Behavioral and Phenotyping System

Promethion Core (Sable Systems, USA) was used for indirect calorimetry experiments. Animals were single caged and acclimatized for two days before experimental procedures were performed. The recording continued for the following seven days. The system consists of a standard GM-500 cage with a food hopper and a water bottle connected to load cells (2 mg precision) with 1 Hz rate data collection. Additionally, the cage contains a red house enrichment. Total (ambulatory and fine) activity was monitored at 1 Hz rate using an XY beam break array (1 cm spacing). Oxygen, carbon dioxide and water vapor were measured using a CGF unit (Sable Systems). This multiplexed system operated in pull-mode. Air flow was measured and controlled by the CGF (Sable Systems) with a set flow rate of 2 L/min. Oxygen consumption and carbon dioxide production were reported in milliliters per minute (mL/min). Energy expenditure (EE) was calculated using the Weir equation ^78^ and Respiratory Exchange Ratio (RER) was calculated as the ratio of VCO_2_/VO_2_. Raw data was processed using Macro Interpreter v2.38 (Sable Systems).

#### Pathogen load quantification

After euthanasia, animals were perfused with ice-cold HBSS and organs were collected to determine pathogen loads. Whole organs were homogenized using a tissue homogenizer. Peritoneal lavage was collected by injecting 5mL of PBS into the peritoneal cavity, followed by shaking the mouse for a few seconds before collecting back the lavage fluid into a syringe. Heparinized blood was obtained by intracardiac puncture. Serial dilutions were plated onto Columbia agar with 5% Sheep Blood (Thermo Fisher Scientific) and pathogen load was calculated by counting Colony-Forming Units (CFUs) after 24-48 h incubation at 37 °C in air 5° CO_2_ (aerobes) or in an air-tight container equipped with the OxoidTM AneroGen™ anaerobe container system (Thermo Fisher Scientific). Anaerobic conditions were confirmed by using an Oxoid ® Resazurin Anaerobic Indicator (Thermo Fisher Scientific).

#### Precision-cut liver slice

Big lobes from livers were embedded in 2.5% low melting Agarose (Carl Roth, 6351.5) and precision cut slices were produced using a vibratome (Compresstome) with the following settings: thickness: 250-300 μM, oscillation: 5.5, speed: 4. Slices were cultured in complete media (DMEM, 10% FBS, L-glutamine, 4.5 g/L glucose, 1% Streptavidin/penicillin) for 1-2 h prior changing into experimental media. To analyze GSK3β signaling, slices were cultured in low-glucose media (DMEM 1.5 g/L glucose, 10% FBS, L-Glutamine, 1% Streptavidin/penicillin) and stimulated with or without LPS (150 ng/mL). Tissues were collected in a time-course manner, snap-frozen, and GSK3β signaling was analyzed by western blot. For glucose output experiments, liver slices were cultured in glucose-free DMEM (Fisher Scientific, Ref: 11520416), supplemented with 20 mM sodium lactate and 2 mM sodium pyruvate. After 24 h incubation, supernatants were collected and the glucose concentration was quantified by using a glucose colorimetric assay kit (Merck, Ref: MAK263-1KT) according to manufacturer’s instructions. Colorimetric signal was normalized to liver slice total protein concentration obtained with the BCA method (Serva).

#### Western Blot

Protein extracts from tissues were obtained by using RIPA buffer (Tris-HCL (10 mM), EDTA (1 mM); Triton X-100 (1%, m/m), sodium deoxycholate (0.1%, m/m), NaCl (140 mM), SDS (0.1%, m/m), protease inhibitor cocktail (Sigma) and phosphatase inhibitor cocktail 3 (Sigma). Protein concentration was quantified by using the BCA method (Serva). 30-50 µg of proteins were loaded and separated in SDS-polyacrilamyde gel electrophoresis under reducing conditions. Proteins were transferred onto a 0.2 mm PVDF membrane using a semi-dry system (Bio-Rad). Membranes were blocked with 5% (w/v) BSA in TBST for 1 h at room temperature and followed by incubation with 1:1000 diluted primary antibodies overnight at 4 °C. After overnight incubation, membranes were washed three times for 5 minutes each with TBST at room temperature (RT) followed by incubation for 2 h at RT with peroxidase-conjugated secondary antibody before detection. Proteins were detected via chemiluminescence by using the SERVALight Eos CL HRP WB Substrate Kit (SERVA) in a Fusion FX Western blot imaging system. Densitometry was performed using ImageJ.

#### Flow cytometry

Single cell suspension from peritoneal lavage was obtained for cell surface staining. Peritoneal lavage was obtained as described in the pathogen load section. Single-cell suspensions were stained in 5 mL polystyrene tubes in FACS buffer (2% BSA + 2 mM EDTA + 2 mM NaN_3_) with the following anti-mouse antibodies to identify distinct immune cell populations: CD45 (clone REA737), CD3 (clone REA641), NK1.1 (clone REA1162), CD11b (clone REA592), Ly-6G (clone REA526), F4/80 (clone REA126), and Ly-6C (clone REA796) (Miltenyi Biotec) for 20 minutes at 4 °C. Cells were stained with propidium iodide (Sigma-Aldrich) for 5 mins at RT after cell surface staining to exclude dead cells in the analysis. For absolute quantification, the CountBright™ Absolute Counting Beads (Invitrogen) kit was used according to the manufacturer’s instructions. Fluorescence minus one (FMO) control was used to identify positive populations in the analysis. Samples were acquired on a BD® LSR II Flow Cytometer and analyzed with FlowJo 10.7.1 software (TreeStar).

#### Cytokines

Cytokines were determined in plasma using the LEGENDplex™ Mouse Inflammation Panel 13-plex (BioLegend, San Diego), according to manufacturer instructions and acquired using a BD Accuri™ C6 Plus Flow Cytometer.

#### qRT-PCR

Livers were collected, snap frozen and kept in −80°C. They were then homogenized with QIAzol lysis reagent (Qiagen, Ref: 79306) using a bead-based tissue lyser (Qiagen, Rf: 85600). Subsequently, chloroform was added, mixed, and incubated for 5 mins at room temperature. The lysate was then transferred into a new tube and centrifuged for 15 mins at 12,000g 4°C. The upper aqueous phase was carefully transferred into a new 1,5 mL reaction tube and fresh isopropanol was added. This mixture was then incubated overnight at −20°C. The samples were centrifuged for 10 min at 12000g, 4°C and the supernatant was carefully removed. The pellet was washed twice with ice-cold 75% ethanol at 7500g 4°C. Pellets were air-dried and dissolved in DEPC-treated water. RNA concentrations were determined with a Nanodrop (ThermoFisher). cDNA was transcribed from total RNA with a M-MuLV First-Strand cDNA Synthesis Kit (NZYTech) following manufactureŕs instructions. Quantitative real-time PCR (qRT-PCR) was performed using 1 µg cDNA and NZYSupreme qPCR Green Master Mix (2x) (NZYTech) in duplicate on a QuantStudio 6 Pro Real-Time PCR System (ThermoFisher Scientific) under the following conditions: polymerase activation (1x cycle, 95°), denaturation and annealing/extension (40x cycles, 95°C and 60°C, respectively). Real-time primer sequences for these samples are in the key resources table. Relative gene expression was calculated using the 2-ΔCT method ^79^ using 60S acidic ribosomal protein P0 (*Rplp0*) as housekeeping control gene.

#### Glycogen measurement

Livers were homogenized with a bead-based tissue lyser (Qiagen, Rf: 85600) in ddH20. Then samples were boiled for 10 mins to inactivate enzymes, followed by centrifugation at 4°C for 10 mins at 12000 g. Supernatants were collected and glycogen was determined by using a colorimetric assay kit (Abcam, Ref: ab65620) using manufacturer’s instructions. The total amount of glycogen was normalized with total proteins quantified via BCA (Serva).

#### Metabolomics

##### Non-targeted metabolomics

Metabolite profiling was conducted using liquid chromatography high-resolution tandem mass spectrometry (LC-HR-MS/MS) following an untargeted metabolomics approach. The instrumentation comprised an ultra-performance liquid chromatography system (A<u>c</u>quity I-class; Waters GmbH, Eschborn, Germany) connected to a quadrupole- time-of-flight mass spectrometer (QToF) equipped with an ionmobility spectrometer (IMS) (VION IMS QToF; Waters GmbH, Eschborn, Germany). For analysis, the sample extract residue was reconstituted in 200 µL aqueous acetonitrile (with 10% water) containing 0.1% formic acid and transferred into autosampler glass vials. Additionally, a pool containing 20 µL from each sample was separately prepared and randomly analyzed multiple times during the analysis process.

Chromatographic separation was achieved by hydrophilic interaction chromatography (HILIC) employing a BEH Amide column (2.1 x 100 mm, 1.7 µm; Waters GmbH, Eschborn, Germany) maintained at 45°C. The mobile phase consisted of a gradient of water including 0.1% formic acid as a mobile phase A and acetonitrile including 0.1% formic acid as a mobile phase B. Ten µL sample volumes were injected at a flow rate of 0.4 mL/min (1%/99% A/B), followed by a linear increase in mobile phase A to 5% within 2 min, reaching the 99% at 7.5 min. After an additional time of 0.5 min, initial conditions were restored within 0.3 min, followed by column equilibration at starting conditions for 1.7 min. For ionization, positive and negative electrospray ionization (ESI) combined with high-definition data acquisition scan mode (HDMSE) was employed, including ion mobility screening to determine metabolite-specific collision cross-section values, accurate mass, and respective fragment ion mass screenings. The observed mass range was set to mass-to-charge ratios between 50 and 1000 Da. The total scan time was set to 0.3 s with 40% of the time applying a collision energy of 6eV (low energy) and the remaining 60% collision energy was ramped from 15 to 50 eV or 10 to 80 eV (high energy) in positive or negative ESI mode, respectively. The instrument parameters for positive ESI mode were set to 1 kV and 40V for capillary voltage and sample cone voltage, including source and desolvation temperatures set at 120°C and 550°C, respectively. In negative ESI mode, capillary voltage and cone voltage were set to 1.5 kV and 40V, including source and desolvation temperatures at 120°C and 450°C, respectively. Nitrogen was used for both cone gas and desolvation gas, with flow rates of 50 L/h and 1000 L/h, respectively, in positive and negative ESI. For mass correction, repeated injections of Leucine-Enkephalin were employed. Data acquisition and raw data processing were conducted using the Waters Unifi software package 2.1.2. Subsequently, Unifi export files derived from positive and negative ESI screenings were imported into the Progenesis QI software (Nonlinear Dynamics, Newcastle upon Tyne, UK) for further data processing. This processing included peak picking with auto threshold and chromatographic alignment using an automatically chosen reference sample, signal deconvolution and normalization of data. For metabolite identification, features from untargeted metabolomics screenings were compared with the human metabolome database (HMDB, www.hmdb.ca) using the Progenesis Metascope plugin, applying a precursor mass accuracy of ≤ 10 ppm and a theoretical fragment mass accuracy of ≤ 10 ppm. Additionally, features were compared to in-house library data achieved by injections of pure metabolite standards in neat solution, including allowed deviations for retention time and CCS of 0.3 min and 3%, respectively. Metabolic features without metabolite annotation were excluded, and the remaining features, along with their signal intensities and respective identifications were exported to Microsoft Excel for further data processing. Metabolomics data were submitted to MetaboAnalyst 6.0 (https://www.metaboanalyst.ca/MetaboAnalyst/ModuleView.xhtml). Metabolites were normalized by median for each sample. One-factor analysis provided principal component analysis (PCA) and selected auto-scaled metabolite concentration values. Quantitative pathway enrichment analysis was based on the Small Molecule Pathway Database (SMPDB) 99 metabolite sets library-based pathway enrichment analysis for significantly regulated metabolites, illustrating enrichment ratio (observed hits / expected hits per pathway) and pathway significance based on individual metabolite p-values.

##### Targeted metabolomics

TCA cycle metabolites and amino acids in livers and muscles from control and septic mice were analysed by using ultrahigh pressure liquid chromatography with tandem mass spectrometry (UHPLC-MS/MS) as described ^80^. Raw data was normalized to tissue weight.

#### Mouse genome scale metabolic modeling

We leveraged genome-scale metabolic modeling to analyze our mouse hypometabolism data. Metabolic modeling was done using the CobraPy Toolbox (v0.26.2) and GUROBI toolbox (v10.0.0) for integer linear programming optimization, while ratio, statistics and visualization were computed using the R programming language. We first downloaded a curated mouse metabolic model (v1.3.1) from https://github.com/SysBioChalmers/Mouse-GEM/releases/tag/v1.3.1 ^40^. Next, uptake rates were parameterized using our metabolomics data. If any measured metabolite was not connected via an exchange (EX) reactions to the model, additional exchange reactions as well as internal transport reactions to the corresponding metabolites were added to the model under the premise that the metabolomics measurements quantify the presence of the data influx allowing for the additional presence of these metabolites (18 in total). Next, we determined a base medium using the minimal_medium function from cobrapy to identify a minimal required medium influx to support at least 95% optimized ATP production. Next, metabolomics ratios were computed for each condition and tissue (liver and muscle) and compared to baseline abundance. Each influx of the minimal base medium was then modified by the calculated ratios per condition. Optimal ATP production was then computed using Flux balance analysis (FBA).

#### Proteomics and Phospho-proteomics

##### Phosphoproteomics and whole proteome sample preparation

For proteomics analysis snap frozen tissues were transferred to Precellys CKMix Lysing Kits tubes (Precellys ®, Bertin Technologies, France, 1.4 mm and 2.8 mm ceramic beads inside) and 1 mL PBS supplemented with Roche cOmplete (Roche) and PhosSTOP (Roche). The tissues were lysed and homogenized using a Precellys 24 (Precellys ®, Bertin Technologies, France) for 3 cycles (20 sec ON/30 sec OFF) at 6000 rpm followed by 30 sec intervals. For further analysis, lysis buffer (final concentration: 1% SDS, 50 mM DTT and 100 mM HEPES pH 8) was added to 100 µg protein and sonicated (Bioruptor Plus, Diagenode, Belgium) for 10 cycles (60 sec ON/30 sec OFF) at high setting and at 20 °C. Then samples were boiled at 95°C for 5 min and alkylated with iodoacetamide (IAA, final concentration 15 mM) for 30 min at room temperature in the dark. Proteins were precipitated overnight at −20 °C after addition of 4x volume of ice-cold acetone. The following day, the samples were centrifuged at 20800 x g for 30 min at 4 °C and the supernatant carefully removed (Eppendorf 5810R, Eppendorf AG, Germany). Pellets were washed twice with ice-cold 80% (^v^/^v^) acetone in water then centrifuged at 20,800x g at 4 °C for 10 min. After removing the acetone, pellets were air-dried before addition of 100 µL of digestion buffer (1 M guanidine chloride in 100 mM HEPES pH 8). Samples were resuspended with sonication as explained above, then LysC (Wako) was added at 1:100 (w/w) enzyme:protein ratio and digestion proceeded for 2 h at 37 °C under shaking (650 rpm). Samples were then diluted 1:1 with MilliQ water and trypsin (Promega) added at 1:100 (^w^/_w_) enzyme:protein ratio. Samples were further digested overnight at 37 °C under shaking (650 rpm). The day after, digests were acidified by the addition of TFA to a final concentration of 1% (^v^/_v_), then desalted with Waters Oasis® HLB µElution Plate 30 µm (Waters Corporation, MA, USA) under a soft vacuum following the manufacturer instruction. Briefly, the columns were conditioned with 3x100 µL solvent B (80% (v/v) acetonitrile; 0.05% (^v^/_v_) formic acid) and equilibrated with 3x100 µL solvent A (0.05% (v/v) formic acid in Milli-Q water). The samples were loaded, washed 3 times with 100 µL solvent A, and then eluted into 0.2 mL PCR tubes with 50 µL OASIS elution buffer (80% ACN and 0.1% TFA buffer solution). Before phosphopeptide enrichment, samples were filled up to 210 µL. Phosphorylated peptides were enriched using Fe(III)-IMAC cartridges (Agilent) in an automated fashion using the standard protocol from the AssayMAP Bravo Platform (Agilent Technologies). In short, Fe(III)-IMAC cartridges were first primed with 100 µL of priming buffer (100% ACN, 0.1% TFA) and equilibrated with 50 μL of OASIS elution buffer. After loading the samples into the cartridge, the cartridges were washed with an OASIS elution buffer, while the syringes were washed with a priming buffer. The phosphopeptides were eluted with 25 μL of 1% ammonia directly into 25 μL of 10% FA. Samples were dried down with a speed vacuum centrifuge and stored at −20 °C until LC-MS analysis. For whole proteome analysis, the flow through after phosphoenrichment was dried down with a speed vacuum centrifuge and stored at −20 °C until LC-MS analysis.

For proteomics analysis of plasma, 5 μL of mouse plasma were treated using ProteoSpin™ Abundant Serum Protein Depletion Kit (Norgen Biotek) according to the manufacturer instruction to deplete the most abundant serum proteins. After elution of the bound proteins, samples were treated with 4 volumes ice cold acetone and left overnight at −20 °C to precipitate the proteins. The samples were then centrifuged at 14000 rpm for 30 minutes, 4 °C. After removal of the supernatant, the precipitates were washed twice with 300 µL of a solution of ice cold 80 % acetone. After addition of each wash solution, the samples were vortexed and centrifuged again for 10 minutes at 4°C. The pellets were then allowed to air-dry before being dissolved in digestion buffer at 1 µg/µL (1M guanidine HCl in 0.1M HEPES, pH 8). To facilitate the resuspension of the protein pellet, the samples were subjected to 5 cycles of sonication in a Bioruptor (Biogenode) with 1 minute on and 30s off with high intensity. Afterwards, LysC (Wako) was added at 1:100 (w/w) enzyme:protein ratio and digestion proceeded for 2 h at 37 °C under shaking (1000 rpm for 1 h, then 650 rpm). The samples were diluted 1:1 with milliQ water (to reach 1.5M urea) and were incubated with a 1:100 w/w amount of trypsin (Promega sequencing grade) overnight at 37 °C, 650 rpm. The digests were then acidified with 10% trifluoroacetic acid and then desalted with Waters Oasis® HLB µElution Plate 30µm in the presence of a slow vacuum. In this process, the columns were conditioned with 3x100 µL solvent B (80% acetonitrile; 0.05% formic acid) and equilibrated with 3x 100 µL solvent A (0.05% formic acid in milliQ water). The samples were loaded, washed 3 times with 100 µL solvent A, and then eluted into PCR tubes with 50 µL solvent B. The eluates were dried using a speed vacuum centrifuge (Eppendorf Concentrator Plus, Eppendorf AG, Germany) and stored at −20° C. Before analysis, samples were reconstituted in in MS Buffer (5% acetonitrile, 95% Milli-Q water, with 0.1% formic acid), spiked with iRT peptides (Biognosys, Switzerland) and loaded on Evotips (Evosep) according to the manufacturer’s instructions. In short, Evotips were first washed with Evosep buffer B (acetonitrile, 0.1% formic acid), conditioned with 100% isopropanol and equilibrated with Evosep buffer A. Afterwards, the samples were loaded on the Evotips and washed with Evosep buffer A. The loaded Evotips were topped up with buffer A and stored until the measurement.

##### LC-MS Data independent analysis (DIA) for whole proteomics and phosphoproteomics

Prior to analysis, samples were reconstituted in MS Buffer (5% acetonitrile, 95% Milli-Q water, with 0.1% formic acid) and spiked with iRT peptides (Biognosys, Switzerland). Peptides from BAT and muscles samples were separated in trap/elute mode using the nanoAcquity MClass Ultra-High Performance Liquid Chromatography system (Waters, Waters Corporation, Milford, MA,USA) while samples from liver were separated in trap/elute mode using a nanoAcquity UPLC (Waters, Milford, MA). Both systems were equipped with a trapping (Waters nanoEase M/Z Symmetry C18, 5 μm, 180 μm x 20 mm) and an analytical column (Waters nanoEase M/Z Peptide C18, 1.7 μm, 75 μm x 250 mm).

For whole proteome analysis, solvent A was water and 0.1% formic acid, and solvent B was acetonitrile and 0.1% formic acid. 1 µL of the sample (∼1 µg on column) were loaded with a constant flow of solvent A at 5 μl/min onto the trapping column. Trapping time was 6 min. Peptides were eluted via the analytical column with a constant flow of 0.3 µL/min. During the elution, the percentage of solvent B increased in a nonlinear fashion from 0-40% in 120 min. Total run time was 145 min. including equilibration and conditioning. For phosphoproteomic analysis, the gradient was shortened: the percentage of solvent B increased in a nonlinear fashion from 0-40% in 60 min. Total run time was 75 min. The LC was either coupled to an Orbitrap Exploris 480 (ThermoFisher Scientific, Bremen, Germany) or to Orbitrap Fusion Lumos (Thermo Fisher Scientific, Bremen, Germany) using the Proxeon nanospray source. The peptides were introduced into the mass spectrometer via a Pico-Tip Emitter 360 μm outer diameter × 20 μm inner diameter, 10 μm tip (New Objective) heated at 300 °C, and a spray voltage of 2.2 kV was applied. The capillary temperature was set at 300°C. The radio frequency ion funnel was set to 30%.

For whole cell analysis on the Orbitrap Fusion Lumos, full scan MS spectra with mass range 350-1650 m/z were acquired in profile mode in the Orbitrap with resolution of 120.000 FWHM. The default charge state was set to 3+. The filling time was set at maximum of 20 ms with an AGC target of 5 x 10^5^ ions. DIA scans were acquired with 34 mass window segments of differing widths across the MS1 mass range. The HCD collision energy was set to 30%. MS/MS scan resolution in the Orbitrap was set to 30,000 FWHM with a fixed first mass of 200m/z after accumulation of 1x 10^6^ ions or after filling time of 70 ms (whichever occurred first). For all data acquisition and processing Tune version 2.1 and Xcalibur 4.1 were employed. The MS/MS scan resolution in the Orbitrap was set to 30k with an AGC target of 1 × 10^6^ and max injection time of 70 ms. For phosphoproteomics, parameters were the same, except for DIA scans that were acquired with 30 mass window segments of differing widths and MS2 filling time was 47 ms.

For analysis on the Exploris 480, full scan mass spectrometry (MS) spectra with mass range 350–1650 m/z were acquired in profile mode in the Orbitrap with resolution of 120,000 FWHM.

The default charge state was set to 3+. The filling time was set at maximum of 60 ms with limitation of 3 × 10^6^ ions. DIA scans were acquired with 40 mass window segments of differing widths across the MS1 mass range. Higher collisional dissociation fragmentation (stepped normalized collision energy; 25, 27.5, and 30%) was applied and MS/MS spectra were acquired with a resolution of 30,000 FWHM with a fixed first mass of 200 m/z after accumulation of 3 × 10^6^ ions or after filling time of 35 ms (whichever occurred first). Data were acquired in profile mode. For data acquisition and processing of the raw data Xcalibur 4.4 (Thermo) and Tune version 3.1 were used. For phosphoproteomics parameters were the same, except: that DIA scans were acquired with 30 mass window segments of differing widths across the MS1 mass range and MS2 filling time was 47 ms.

For plasma analysis and some liver samples, peptides were separated using the Evosep One system (Evosep, Odense, Denmark) equipped with a 15 cm x 150 μm i.d. packed with a 1.5 μm Reprosil-Pur C18 bead column (Evosep performance, EV-1137, Denmark). The samples were run with a pre-programmed proprietary Evosep gradient of 44 min (30 samples per day) using water and 0.1% formic acid and solvent B acetonitrile and 0.1% formic acid as solvents. The LC was coupled to an Orbitrap Exploris 480 (Thermo Fisher Scientific, Bremen, Germany) using PepSep Sprayers and a Proxeon nanospray source. The peptides were introduced into the mass spectrometer via a PepSep Emitter 360-μm outer diameter × 20-μm inner diameter, heated at 300°C, and a spray voltage of 2 kV was applied. The injection capillary temperature was set at 300°C. The radio frequency ion funnel was set to 30%. For DIA data acquisition, full scan mass spectrometry (MS) spectra with a mass range of 350–1650 m/z were acquired in profile mode in the Orbitrap with a resolution of 120,000 FWHM. The default charge state was set to 2+, and the filling time was set at a maximum of 20 ms with a limitation of 3 × 106 ions. DIA scans were acquired with 40 mass window segments of differing widths across the MS1 mass range. Higher collisional dissociation fragmentation (normalized collision energy 30%) was applied, and MS/MS spectra were acquired with a resolution of 30,000 FWHM with a fixed first mass of 200 m/z after accumulation of 1 × 106 ions or after filling time of 45 ms (whichever occurred first). Data were acquired in profile mode. For data acquisition and processing of the raw data, Xcalibur 4.4 (Thermo) and Tune version 4.0 were used.

##### Data processing

DIA raw data were analyzed using the directDIA pipeline in Spectronaut (v.15 to v. 20 depending on the experiment, Biognosysis, Switzerland). The data were searched against a species specific (*Mus Musculus*, 16.747 entries) and a contaminant (247 entries) Swissprot database.

For whole proteome analysis, the data were searched with the following variable modifications: Oxidation (M) and Acetyl (Protein N-term). A maximum of two missed cleavages for trypsin and five variable modifications were allowed. The identifications were filtered to satisfy FDR of 1 % on peptide and protein level. Relative quantification was performed in Spectronaut for each paired comparison using the replicate samples from each condition. The data (candidate table) and data reports (protein quantities) were then exported and further data analyses and visualization were performed with Rstudio using in-house pipelines and scripts. To select significant proteins, a log_2_FC cutoff of 0.58 and a q-value <0.05 were defined.

For phosphoproteomics analysis, the data were searched with the following modifications: Carbamidomethyl (C) (Fixed) and Oxidation (M), Acetyl (Protein N-term), Phospho (STY) (Variable). PTM localization probability was set to 0.75 and consolidation of phosphosites was sum based. A maximum of 2 missed cleavages for trypsin and 5 variable modifications were allowed. The identifications were filtered to satisfy FDR of 1 % on peptide and protein level. Relative quantification at phospho-site level was performed in Spectronaut for each paired comparison using the replicate samples from each condition. The data (candidate table) and data reports (protein quantities) were then exported and further data analyses and visualization were performed with Rstudio using in-house pipelines and scripts. To select significant phosphosites, a log_2_FC cutoff of 0.58 and a q-value <0.05 were defined. For identifying the most regulated kinases and the corresponding regulated top phosphosites, a modified version of PhosR ^44^ was used.

#### Human analysis

To analyse the association of our murine results to human data we first mapped the identifiers of our murine data to human orthologues using g:profiler’s conversion functionality (https://biit.cs.ut.ee/gprofiler/convert). We augmented this list for blastp search of human orthologues, which ultimately resulted in a list of 118 orthologues human protein IDs. We next investigated statistics and fold changes reported in human plasma proteomics from three different sources. Specifically, we downloaded the data file S3 from Mi et al. 2024 ^59^ (url: https://www.science.org/doi/suppl/10.1126/scitranslmed.adh0185/suppl_file/scitranslmed.adh0185_data_files_s1_to_s3.zip) Supplementary Table S3 from Liang et al. 2021 (url: https://www.life-science-alliance.org/content/lsa/4/10/e202101091/DC3/embed/inline-supplementary-material-3.xlsx) as well as Supplementary Table S2 and S7 from Palmowski et al., 2025 ^60^ (url: https://www.thelancet.com/cms/10.1016/j.ebiom.2024.105508/attachment/bdc81cec-94ce-4d48-a042-fe3cbc296f30/mmc1.docx). We cleaned and filtered these datasets for our orthologues candidate IDs and visualised their comparison to our murine data using the R programming language.

#### Acutelines human cohort

Data used for *Fig. 4I* was derived from the Acutelines data- and biobank, a prospective data, image, and biobank at the emergency department (ED) of the University Medical Center Groningen (UMCG), University of Groningen, the Netherlands (METC#2019/589; ClinicalTrials.gov NCT04615065). Patients aged ≥18 years presenting to the ED with a clinical suspicion of infection were included. Infection status was confirmed through post-hoc clinical adjudication by two independent physicians. Severe outcome was defined as ICU admission and/or death within 7 days of ED presentation. A deferred consent procedure was in place; where proxy consent could not be obtained, an opt-out procedure was followed. Patient characteristics are provided in *Table 1*.

Plasma proteomics was performed using the Olink® Explore HT platform (Olink Proteomics, Uppsala, Sweden), which simultaneously quantifies 5,416 protein assays via proximity extension assay (PEA) technology. Briefly, pairs of oligonucleotide-labeled antibodies bind to target proteins in plasma; the resulting unique hybridization sequences are amplified and quantified by real-time PCR. Protein abundances are reported on a relative log_2_ scale as Normalized Protein eXpression (NPX) values. All samples passed the platform’s internal quality controls.

#### Statistical analyses

Survival is presented as Kaplan-Meier plots and were analyzed using the log-rank test. D’Agostino-Pearson and Shapiro-Wilk normality tests were used to determine normality distribution on data sets. Comparison between two-independent groups was performed by using unpaired two-tailed Student’s t-test or Mann-Whitney test. Comparison between two-dependent groups was performed by paired two-tailed Student’s t-test or Wilcoxon matched-pair test. When more than two groups were compared, one-way analysis of variance (ANOVA), Krurskal-Wallis test followed by Tukey post hoc test. Two-way ANOVA with Tukey multiple comparison was performed for time course experiments. The specific statistical test used is depicted in the figure legends. Data are presented as mean ± SD when appropriate for independent experiments as indicated in each figure legend. Pearson’s correlation coefficient (r) was used for correlation analysis, and 95% confidence intervals are shaded. Experiments were performed with biologically independent animals for each condition and experiments were repeated at least two times as indicated in the figure legends. We used both male and female mice and allocation was randomized. Blinding was performed in experiments using genetically modified mice strains; however, it was not possible for infection experiments comparing to non-infected mice, since the outcome was readily apparent to investigators. Likewise, experiments involving GSK3β inhibition was not blinded due to specific color of the solubilized inhibitor. Significance was accepted when *P* value was below 0.05 and depicted as: *< 0.05, **<0.01, ***<0.001, ****<0.0001. Statistical analysis was performed using GraphPad Prism version 10.6.1 (GraphPad Software Inc., USA).

## SUPPLEMENTAL FIGURES

**Figure S1.**
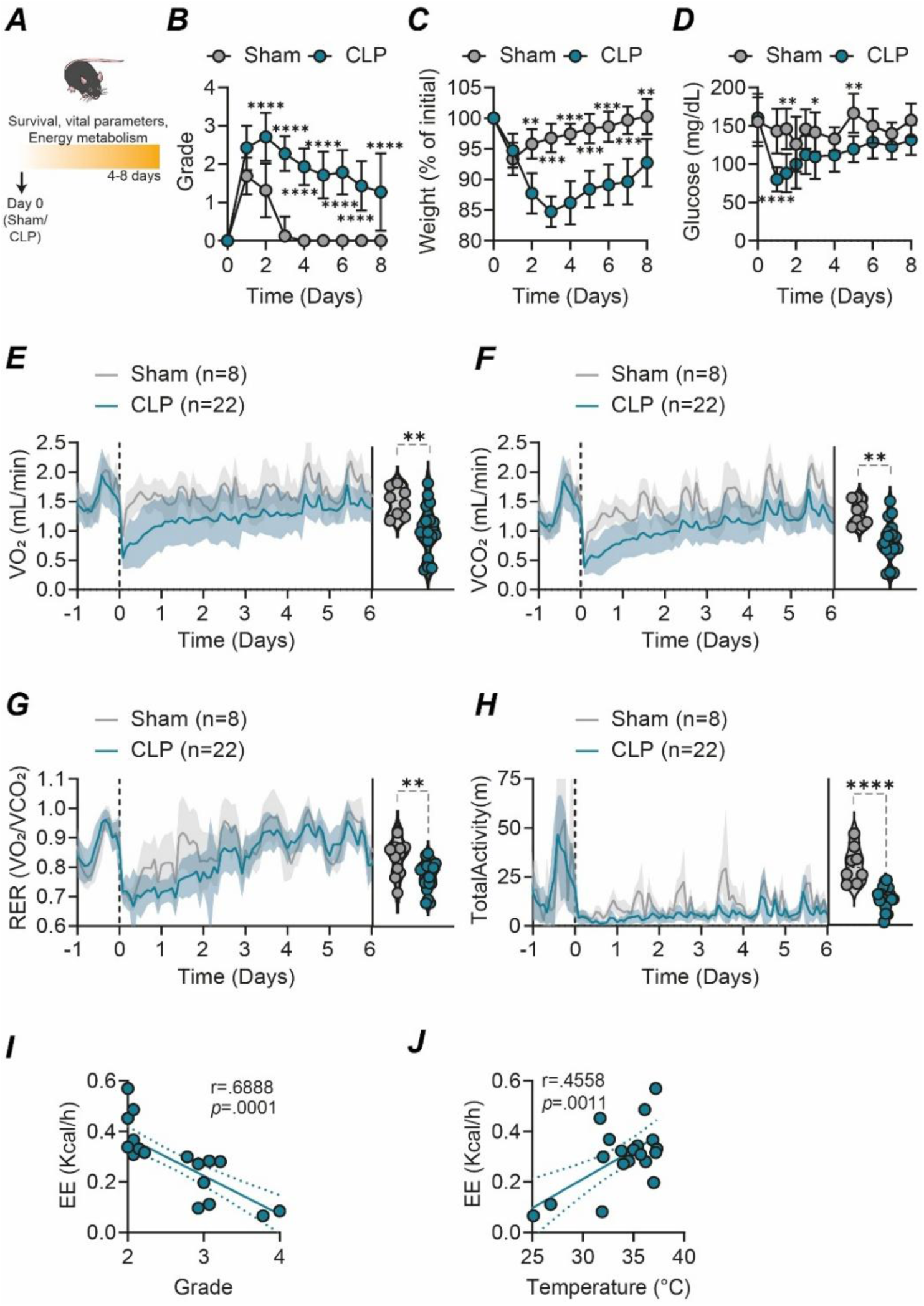
Polymicrobial sepsis triggers a conserved hypometabolic response. Related to Fig. 1. **(A)** Experimental setup. **(B)** Clinical severity score (Grade), **(C)** relative weight and **(D)** blood glucose levels from C57Bl/6j mice that were subjected to either a Sham (n=12) or polymicrobial sepsis via a cecal-ligation and puncture (CLP, n=22), pooled from 6 independent experiments. **(E)** VO_2_, **(F)** VCO_2_, **(G)** respiratory exchange ratio (RER, VCO_2_/VO_2_) and **(H)** total activity from Sham (n=12) or CLP (n=20) mice. The start of the surgery is depicted in a vertical dotted black line in time 0h. Line in the curve indicates the mean, and shading denotes ± SD. Violin plot depicts the mean data from individual mice throughout the complete duration of the experiment. Data pooled from 6 independent experiments **(I)** Correlation between EE vs clinical severity score from mice subjected to CLP. **(J)** Correlation between EE vs body temperature from mice subjected to CLP. Survival was represented by Kaplan-Meier plots and the difference between groups was assessed using the log-rank test. Differences between two groups was assessed with a two-tailed T test. Pearson’s correlation coefficient (r) was used for correlation analysis and 95% confidence intervals are shaded. Data was obtained from at least three independent experiments. *p < 0.05; **p < 0.01; ***p < 0.001; ****p < 0.0001. <u>Abbreviations:</u> m.. meter; RER.. respiratory exchange ratio; VCO_2_.. carbon dioxide production; VO_2_.. oxygen consumption.

**Figure S2.**
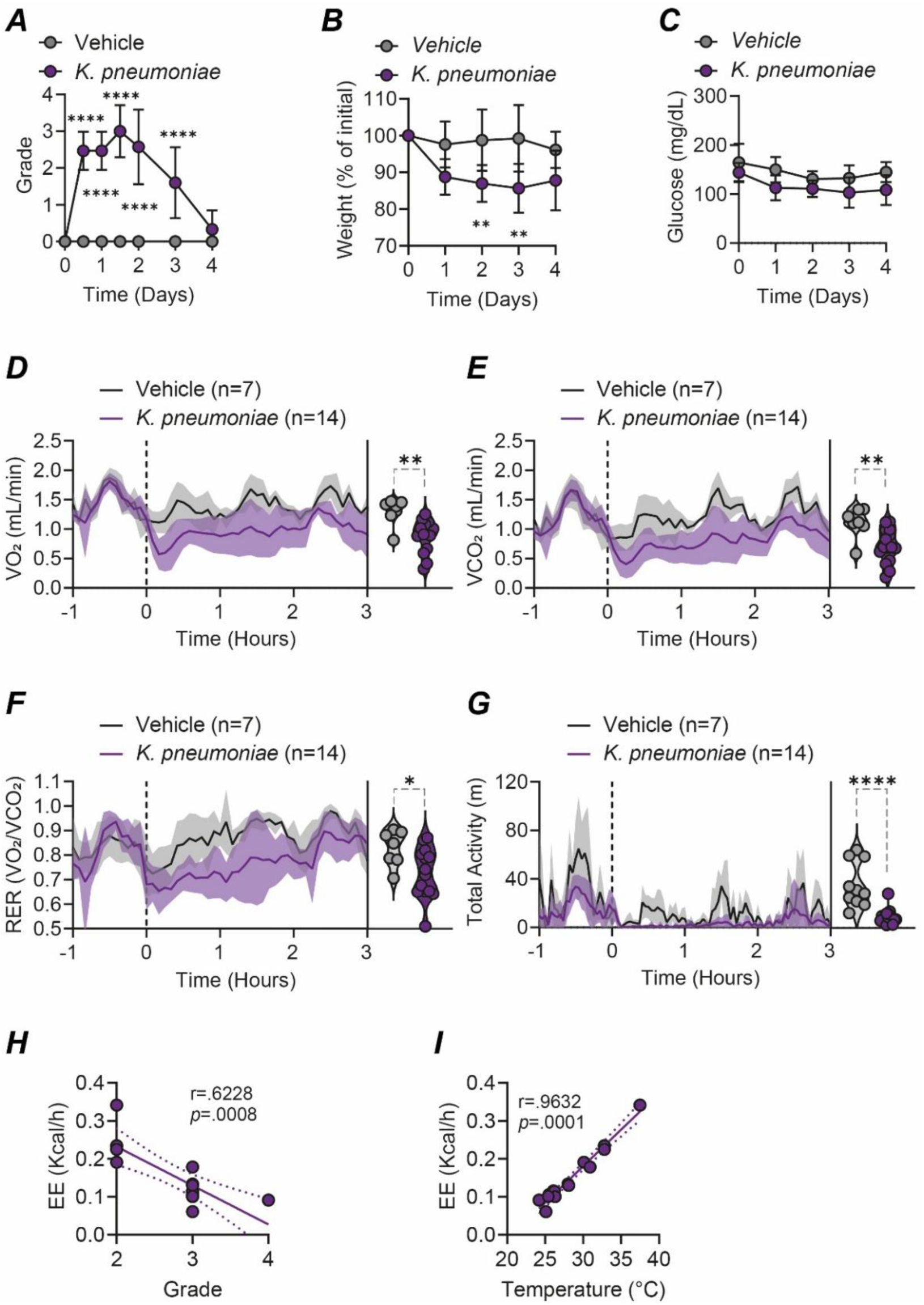
Severe pulmonary infection with K.pneumoniae triggers a conserved hypometabolic response. Related to Fig. 1. (**A**) Disease severity (Grade), (**B**) relative weight, (**C**) blood glucose, (**D**) VO2, (**E**) VCO2, (**F**) RER, and (**G**) total activity from mice subjected to either vehicle (n=7) or lung infection with *K.pneumoniae* (n=19). Data pooled from 4 independent experiments. **(H)** Correlation between EE *vs.* clinical severity score from mice subjected to *K.pneumoniae*. **(I)** Correlation between EE *vs.* body temperature from mice subjected to *K.pneumoniae*. Survival was represented by Kaplan-Meier plots and the difference between groups was assessed using the log-rank test. Differences between two groups was assessed with a two-tailed T test. Pearson’s correlation coefficient (r) was used for correlation analysis and 95% confidence intervals are shaded. *p < 0.05; **p < 0.01; ***p < 0.001; ****p < 0.0001. <u>Abbreviations:</u> m.. meter; RER.. respiratory exchange ratio; VCO_2_.. carbon dioxide production; VO_2_.. oxygen consumption.

**Figure S3.**
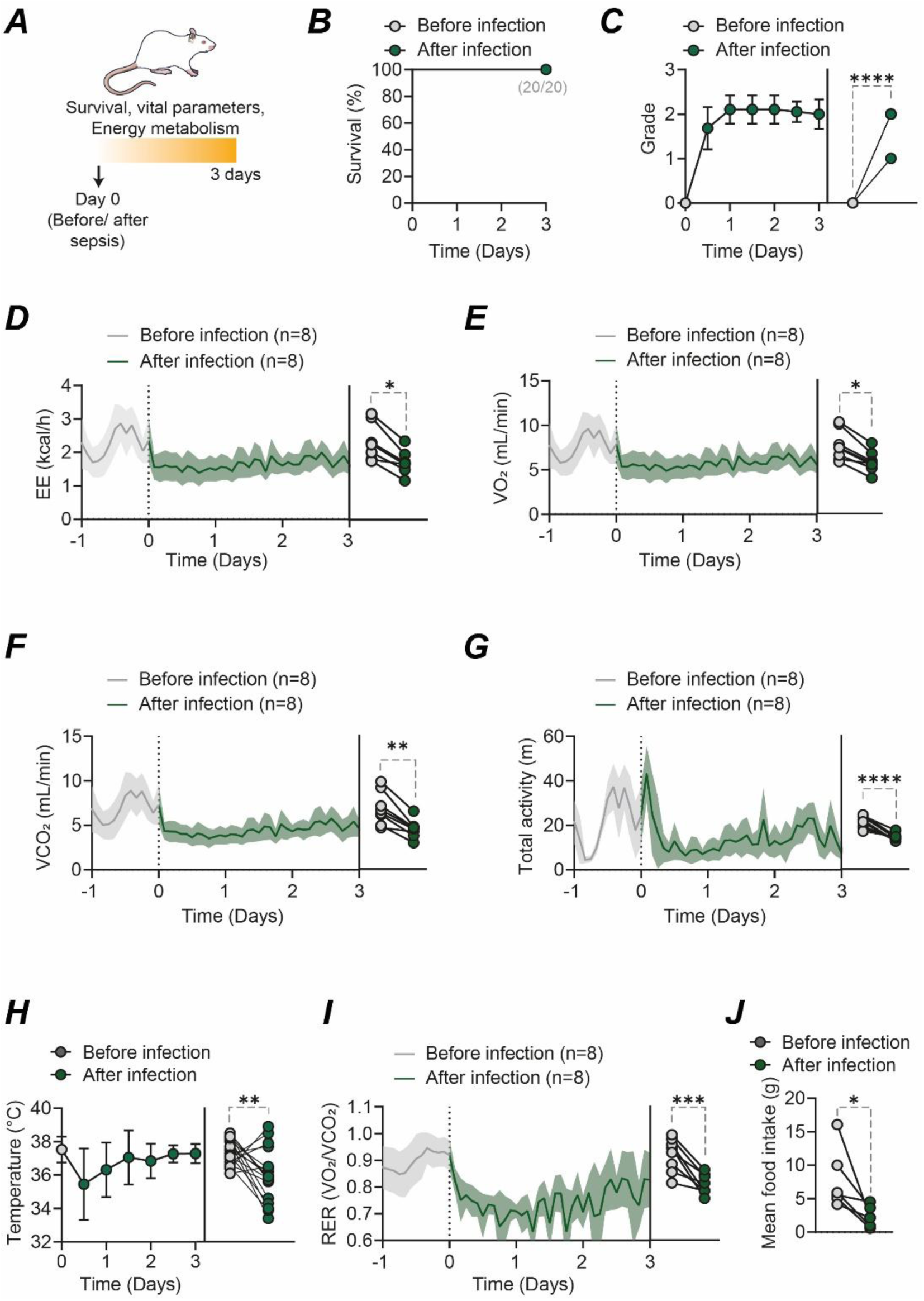
Severe polymicrobial sepsis triggers a conserved hypometabolic response in rats. Related to Fig. 1**. (A)** Experimental setup. **(B)** Survival, **(C)** Clinical severity score (CSS), **(D)** EE, **(E)** VO_2_, **(F)** VCO_2_, **(G)** total activity (n=6), **(H)** body temperature, **(I)** RER, and **(J)** mean food intake before and after rats were subjected to septic peritonitis (n=6-20). The start of the surgery is depicted in a vertical dotted black line in time 0 h. Line in the curves indicates the mean, and shading denotes ± SD. Data from **A**, **B** and **G** is pooled from 6 independent experiments. Data from **C**, **D**, **E**, **F**, **H** and **I** is pooled from 2 independent experiments. Paired comparison depicts the mean data from individual mice before and after infection. Survival was represented by Kaplan-Meier plots and the difference between groups was assessed using the log-rank test. Differences between two groups was assessed with a paired two-tailed T test. *p < 0.05; **p < 0.01; ***p < 0.001; ****p < 0.0001. <u>Abbreviations:</u> m.. meter; RER.. respiratory exchange ratio; VCO_2_.. carbon dioxide production; VO_2_.. oxygen consumption.

**Figure S4.**
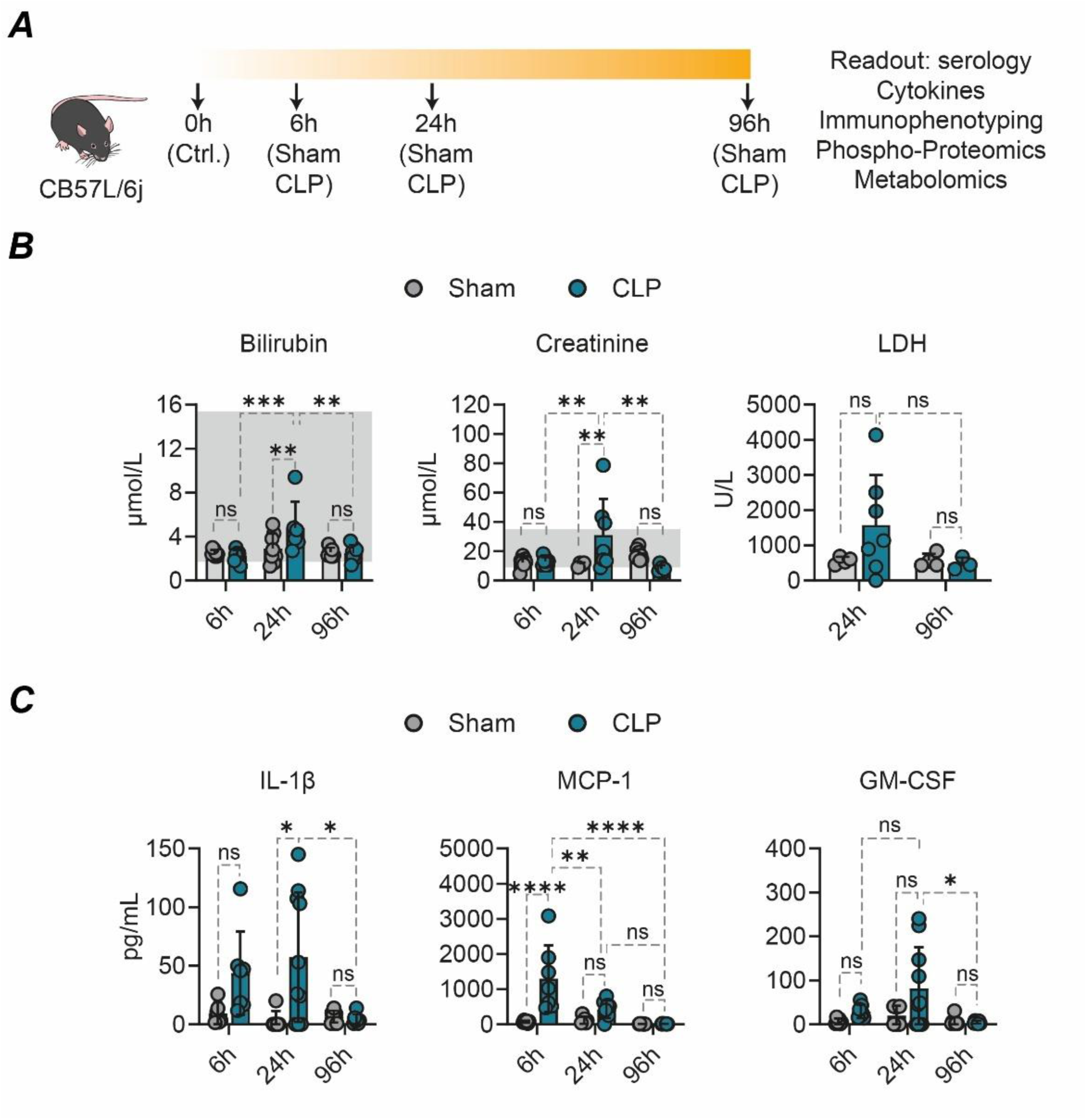
Tissue damage markers and inflammation during polymicrobial sepsis. Related to Fig. 1. **(A)** Animals were subjected to either sham or CLP and organs were collected at different time points depicting the different phases of septic hypometabolism: 6h, 24h (torpor-like), and 96h (recovery). **(B)** Serological markers from mice subjected to sham (n=20) or CLP (n=23) during the torpor-like (6h and 24h) and recovery (96h) phases. Grey bar in the background depict the reference value in healthy mice. **(C)** Cytokine levels in plasma from mice subjected to sham (n=22) or CLP (n=24) during the torpor-like (6h and 24h) and recovery (96 h) phases. Bars indicate mean ± SD and each point represents data from one individual mouse from 4 independent experiments. Comparison between groups were performed by using a Two-way ANOVA with post hoc Tukey test. ns: non-significant, *p < 0.05; **p < 0.01; ***p < 0.001. <u>Abbreviations</u>: GM-CSF.. granulocyte-macrophage colony-stimulating factor; IL.. interleukin; LDH.. Lactate dehydrogenase; MCP-1.. monocyte chemoattractant protein-1.

**Figure S5.**
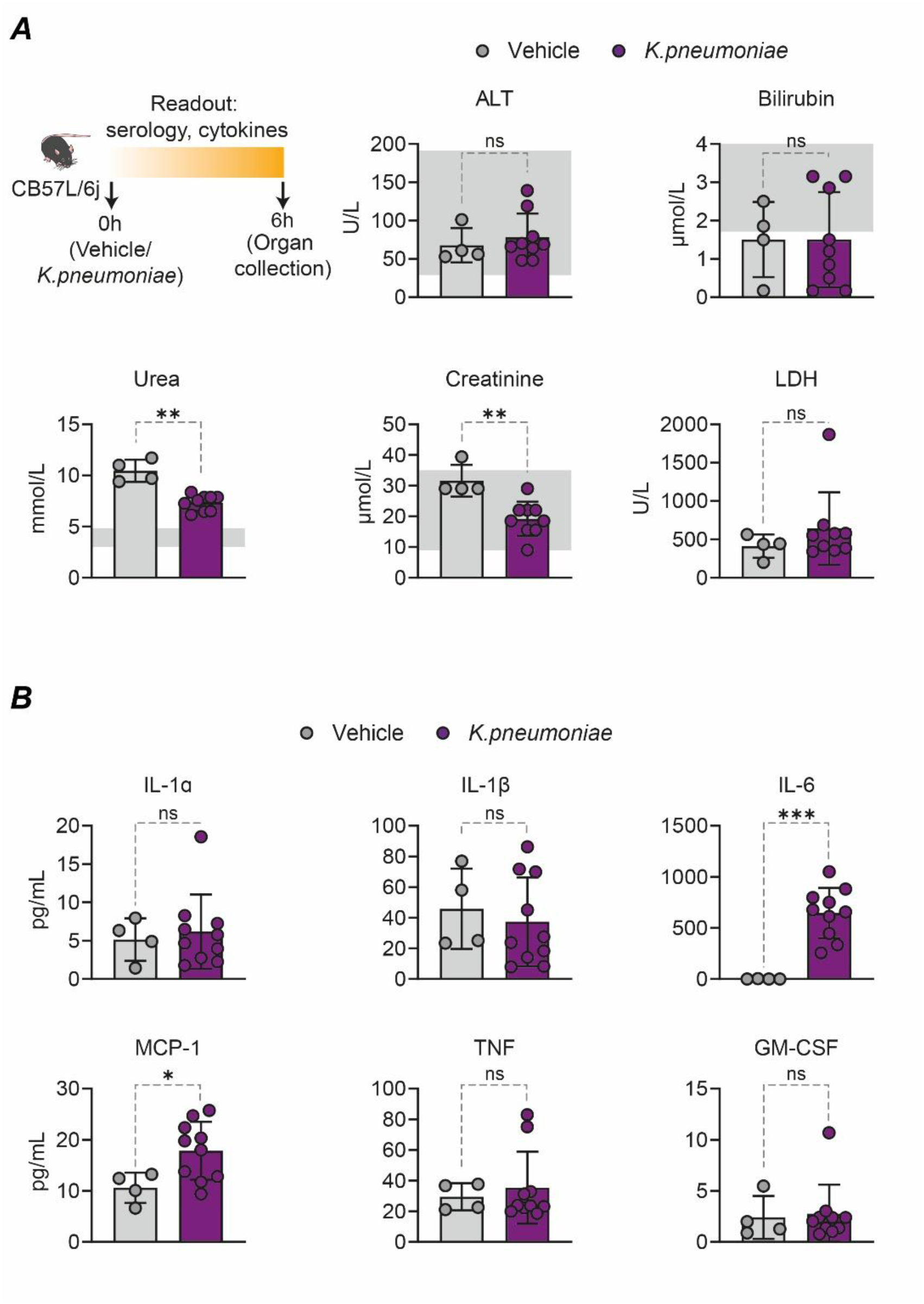
Tissue damage markers and inflammation in the mice subjected to *K.pneumoniae* infection. Related to Fig.1. **(A)** Serological markers 6 h after young mice were subjected to vehicle (n=4) or *K.pneumoniae* infection (n=10). Grey bar in the background depicts the reference value in healthy mice. **(B)** Cytokine levels 6 h after young mice were subjected to vehicle (n=4) or *K.pneumoniae* infection (n=10). Bars indicate mean +/- SD and each point represents data from one individual mouse from 2 independent experiments. Differences between two groups was assessed with an unpaired two-tailed T test. ns: non-significant; *p < 0.05; **p < 0.01; ***p < 0.001. <u>Abbreviations:</u> ALT.. alanine aminotransferase; GM-CSF.. granulocyte-macrophage colony-stimulating factor; IL.. interleukin; LDH.. Lactate dehydrogenase; MCP-1…..monocyte chemoattractant protein-1; TNF.. Tumor necrosis factor.

**Figure S6.**
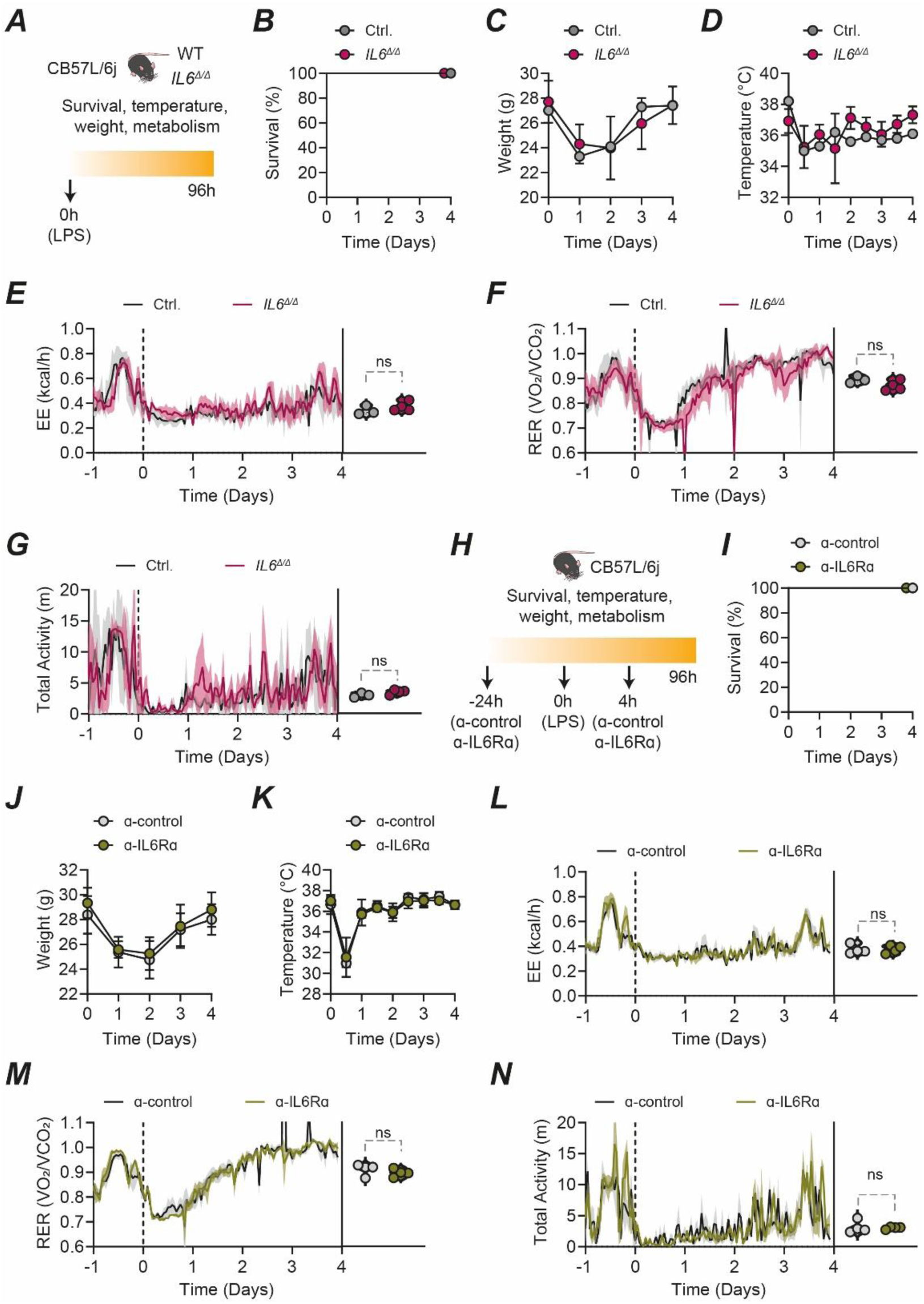
IL-6 does not play a role in sickness behavior and hypometabolism to systemic inflammation. Related to Fig. 1. **(A)** Wild-type or *Il6^-/-^* were subjected to an *i.p.* injection of LPS (5 mg/kg body weight). **(B)** Survival, **(C)** body weight, **(D)** body temperature, **(E)** EE, **(F)** RER, and **(G)** total activity of either wild-type (n=3) or *Il6^Δ/Δ^* (n=4) mice subjected to LPS. **(H)** Wild-type mice were subjected 24 h prior to LPS *i.p.* injection to either a control antibody (α-control) or a blocking antibody against the receptor alpha for IL6 (IL6Rα). Injection of antibodies was repeated 4 h post LPS injection, and the survival, vital parameters, and organismal metabolism was assessed for 96h. **(I)** Survival, **(J)** body weight, **(K)** body temperature, **(L)** EE, **(M)** RER, and **(N)** total activity of mice subjected with either an antibody against a control peptide (n=4) or anti-IL6Ra (n=4) followed by an LPS injection (5mg/kg body weight). Violin plots in **E**, **F**, **G**, **L**, **M** and **N** represents the mean data for the duration of the experiment and each point is data from one individual mouse from 2 independent experiments. ns: non-significant. <u>Abbreviations:</u> EE.. energy expenditure; IL.. interleukin; RER.. respiratory exchange ratio; m.. meters.

**Figure S7.**
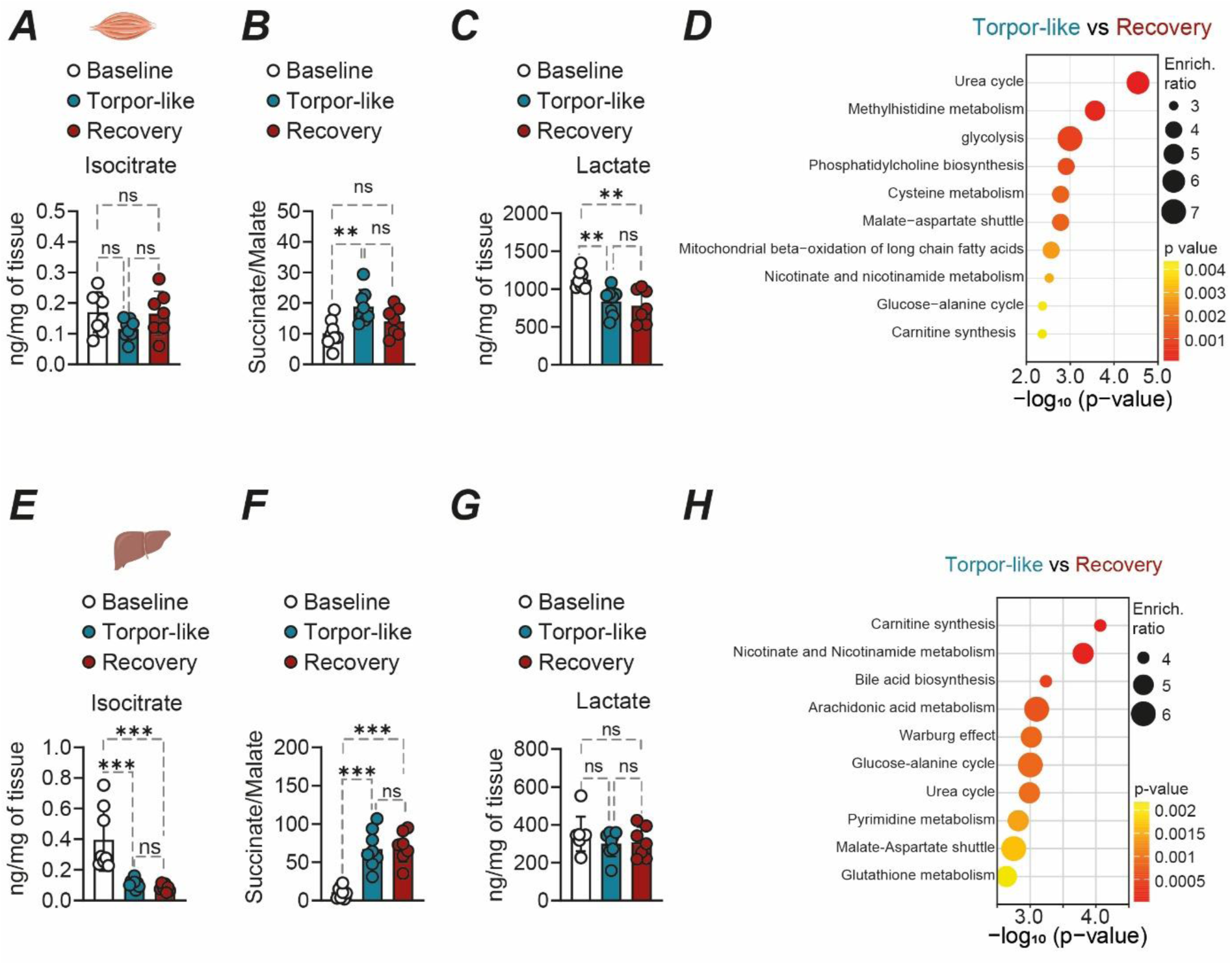
Targeted and untargeted metabolomics at baseline, torpor-like and recovery phases. Related to Fig. 2. **(A)** Isocitrate level, **(B)** Succinate/malate ratio and **(C)** Lactate level in skeletal muscle from mice at the baseline (n=8), torpor-like (n=8) and recovery (n=8) phases. **(D)** SMPDB library-based pathway enrichment analysis for significantly regulated metabolites in skeletal muscle from mice at the torpor-like and recovery phases. **(E)** Isocitrate level, **(F)** Succinate/malate ratio and **(G)** Lactate level in liver from mice at the baseline (n=8), torpor-like (n=8) and recovery (n=8) phases. **(H)** SMPDB library-based pathway enrichment analysis for significantly regulated metabolites in liver from mice at the torpor-like and recovery phases. Comparison between groups was performed by using one-way ANOVA with post hoc Tukey test. **p < 0.01; ***p < 0.001. <u>Abbreviations</u>: ns.. non-signifcant.

**Figure S8.**
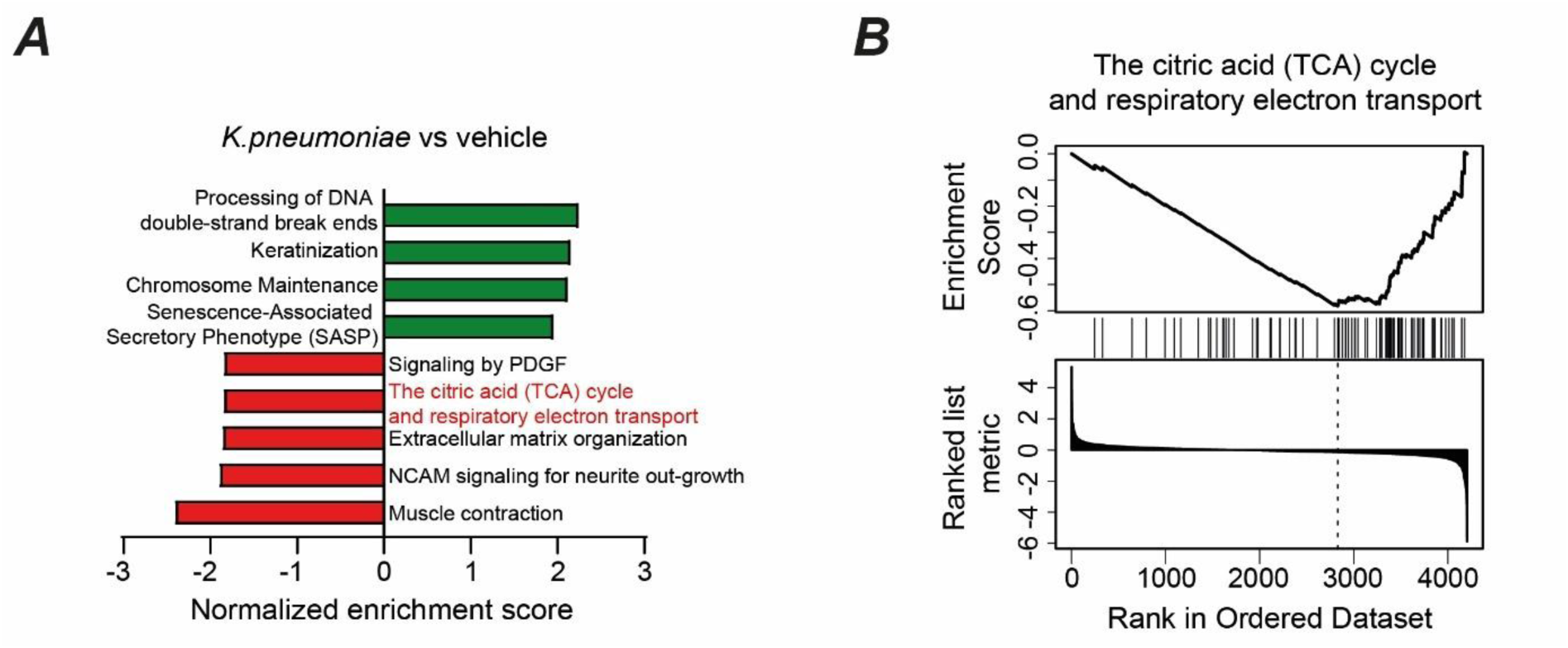
Liver proteomics from mice subjected to *K.pneumoniae* infection. Related to Fig. 2**. (A)** GSEA analyses from liver of mice infected with *K.pneumoniae* compared to mice subjected to vehicle. **(B)** GSEA plot of the citric acid (TCA) cycle enriched in *K.pneumoniae* infected mice compared to vehicle treated mice.

**Figure S9.**
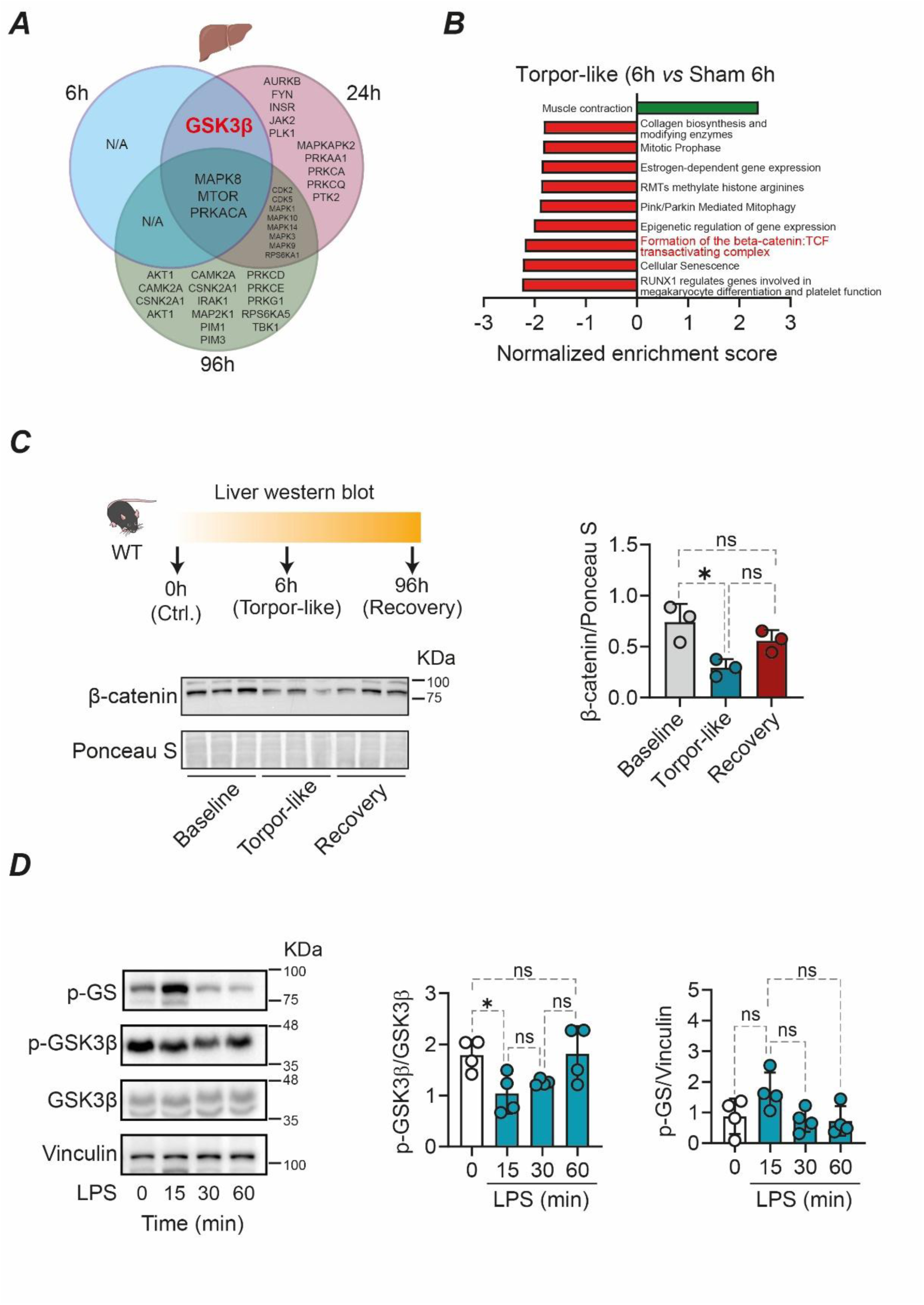
GSK3β activation in the liver. Related to Fig. 3**. (A)** Venn diagram depicting the predicted upregulated kinases in livers obtained at different time points from mice subjected to CLP and normalized to Sham. **(B)** Gene-set enrichment analysis from livers obtained from CLP (n=8) and sham (n=8) at the torpor-like (6h) phase. **(C)** Western blot from liver of mice at baseline, torpor-like and recovery phases. Each lane was loaded from proteins obtained from independent animals. Statistical analysis is show in the right panel. **(D)** Western blot analyzing the phosphorylation of GSK3β and glycogen synthase (GS) from liver slices stimulated with LPS. Densitometric analysis is shown in the right panel. Comparison between groups was performed by using a one-way ANOVA with post hoc Tukey test. *p < 0.05. <u>Abbreviations:</u> CLP..cecal ligation and puncture; GS.. glycogen synthase.

**Figure S10.**
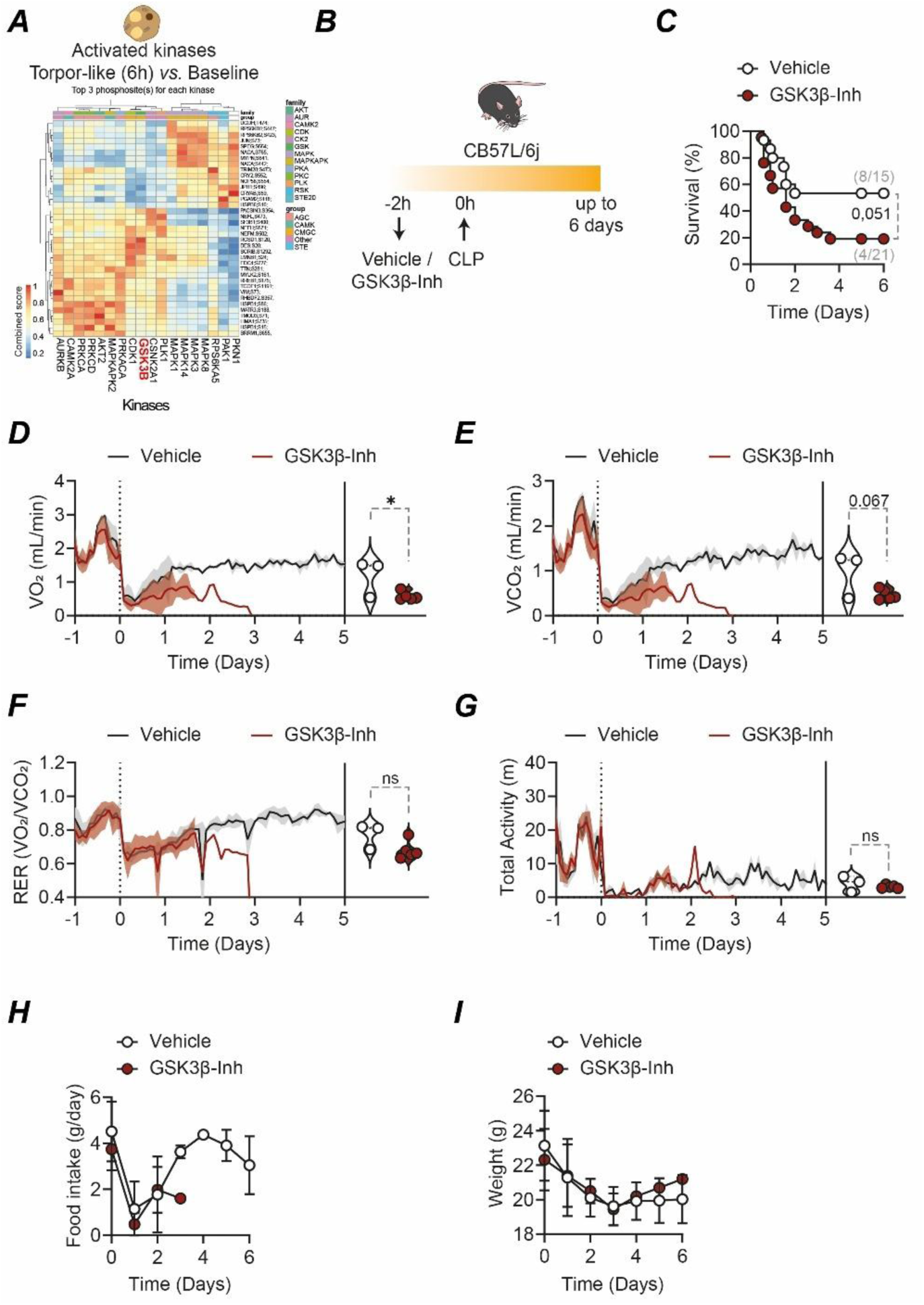
GSK3β activation is necessary to establish disease tolerance to infection. Related to Fig. 3. **(A)** Clustered heatmap of the combined kinase-substrate score for the top three phosphosites of all evaluated kinases. The predicted kinases are depicted at the bottom of the heatmap. Analysis was performed from datasets obtained from brown adipose tissue (BAT) comparing CLP (n=8) vs control (n=8) mice at the 6 h time-point. **(B)** Experimental setup. **(C)** Survival from mice injected with vehicle (n=15) or GSK3β inhibitor (SB216763;25mg/kg body weight) (n=21) 2 h prior CLP. Data pooled from 3 independent experiments. **(D)** VO_2_, **(E)** VCO_2_, **(F)** RER, and **(G)** total activity from mice subjected to CLP and either 2h prior with vehicle (n=3) or GSK3β inhibitor (SB216763) (n=5). The start of the surgery is depicted in a vertical dotted black line in time 0h. Line in the curve indicates the mean, and shading denotes ± SD. Violin plots depict the mean data from individual mice throughout the complete duration of the experiment from 2 independent experiments. **(H)** daily food intake and **(I)** body weight from mice subjected to CLP and either vehicle (n=8) or GSK3β inhibitor (n=12). Survival was represented by Kaplan-Meier plots and the difference between groups was assessed using the log-rank test. Comparison between two groups in **D** was performed using a Mann-Whitney test. Differences between two groups in **E**, **F** and **G** was assessed with an unpaired two-tailed T test. ns: non-significant, *p < 0.05; **p < 0.01. <u>Abbreviations:</u> GSK3β… glycogen synthase kinase 3 beta; CLP… cecal ligation and puncture; m… meters; RER… respiratory exchange ratio; VCO_2_… carbon dioxide production; VO_2_… oxygen consumption.

**Figure S11.**
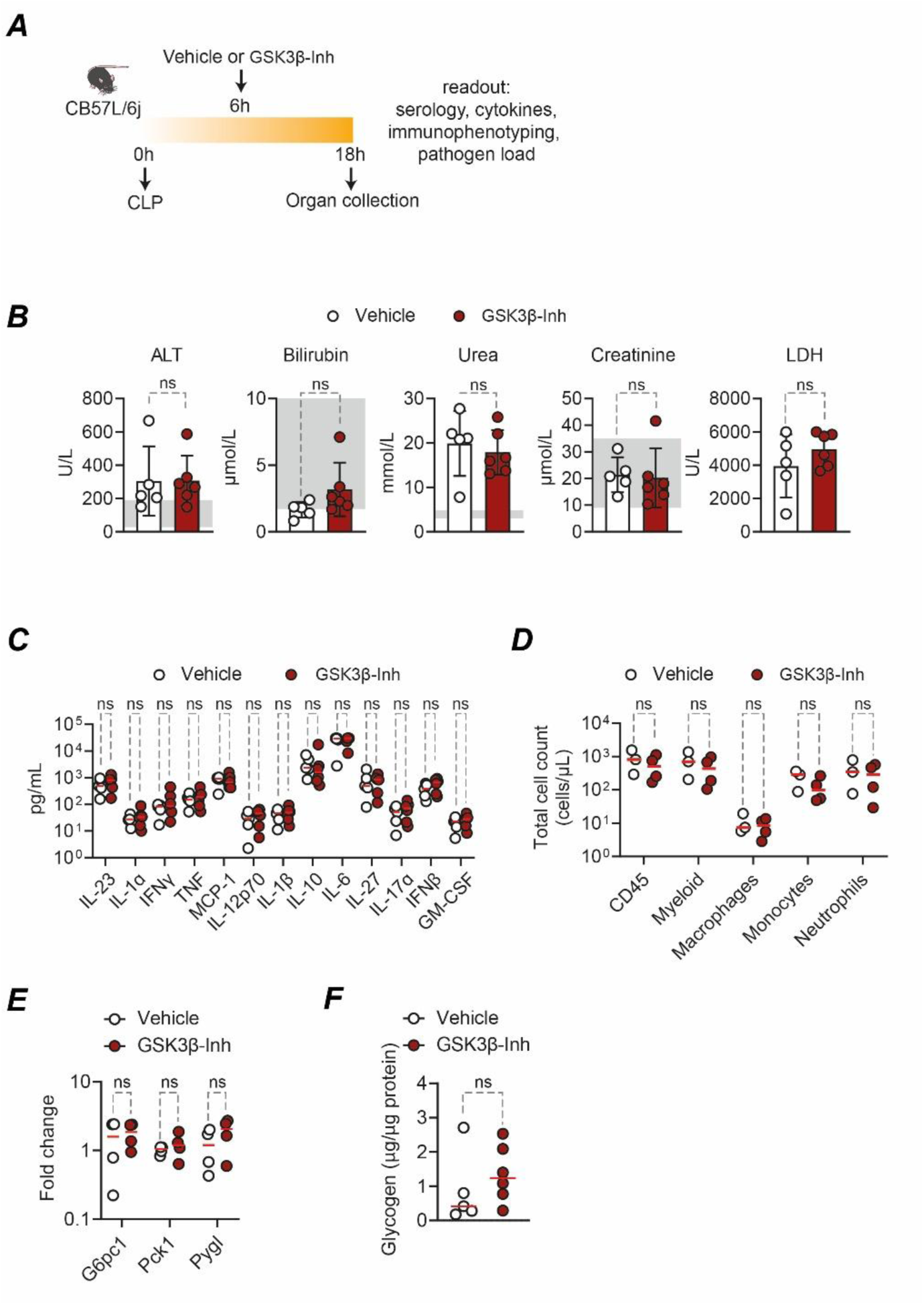
GSK3β inhibition does not increase tissue damage or modulate resistance mechanisms during polymicrobial sepsis. Related to Fig. 3. (**A**) Experimental setup for organ collection of CLP mice subjected to either vehicle or GSK3β inhibitor. (**B**) Serological markers for tissue damage from mice subjected to CLP and either vehicle (n=5) or a GSK3β inhibitor (n=6). Bars denote mean ± SD and each point represent data from one individual mouse from 2 independent experiments. Grey background depicts the reference values from baseline mice. (**C**) Cytokines levels in plasma and (**D**) immune cell frequencies in the peritoneum and from mice subjected to CLP and either vehicle (n=3-5) or a GSK3β inhibitor (n=4-6). **(E)** mRNA levels in the liver of gluconeogenic and glycogenolytic genes from CLP mice subjected to vehicle (n=4) or GSK3β inhibitor (n=4). Data was compared with unpaired T test. (**F**) Glycogen levels of livers from CLP mice subjected to either GSK3β inhibitor (n=5) or vehicle (n=6). Red lines denote the mean. each point represents data from one individual mouse from 2 independent experiments. Comparison between groups was assessed with an unpaired two-tailed T test. ns: non-significant. <u>Abbreviations:</u> ALT… alanine aminotransferase; CLP… cecal ligation and puncture; LDH… Lactate dehydrogenase; IL… interleukin; GM-CSF… granulocyte-macrophage colony-stimulating factor; G6pc1… glucose 6-phosphatase 1; MCP-1… monocyte chemoattractant protein-1; Pck1… phosphoenolpyruvate carboxykinase 1; Pygl… glycogen phosphorylase; TNF… Tumor necrosis factor.

**Figure S12.**
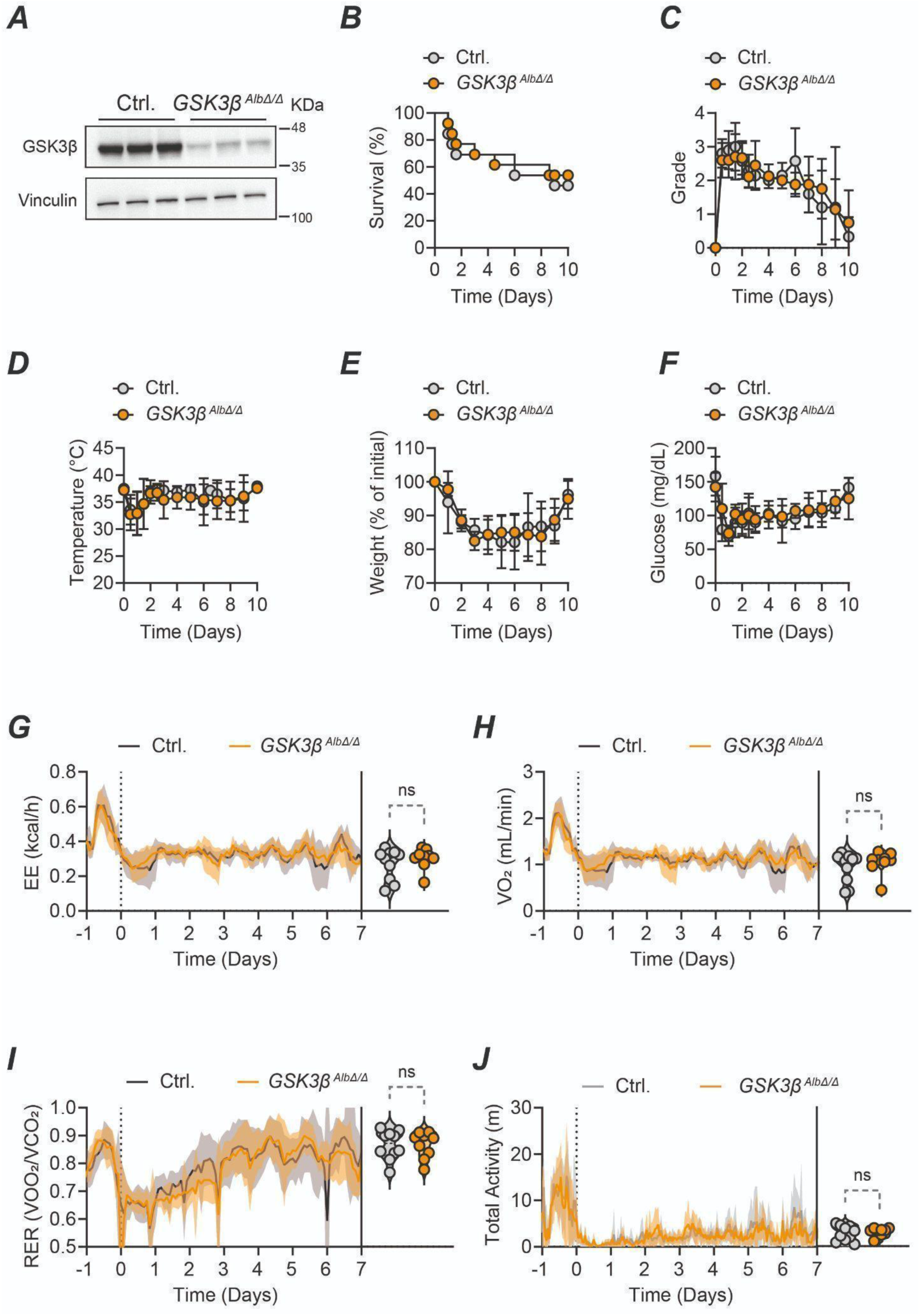
GSK3β in hepatocytes does not control disease severity and metabolism during CLP. Related to Fig. 3**. (A)** Western blot analysis of liver GSK3β expression from control (n=3) and *Gsk3b^AlbΔ/Δ^* mice (GSK3βKO, n=3). **(B)** survival, **(C)** clinical severity score, **(D)** body temperature, **(E)** weight, and **(F)** blood glucose levels from control and *Gsk3b^AlbΔ/Δ^* subjected to CLP. Data is shown as mean ± SD from data obtained in 4 independent experiments. **(G)** EE, **(H)** VO_2_, **(I)** VCO_2_, and **(J)** RER were measured from control and *Gsk3b^AlbΔ/Δ^* mice subjected to CLP. The start of the surgery is depicted in a vertical dotted black line in time 0h. Line in curves indicates the mean, and shading denotes ± SD. Violin plots depict the mean data from individual mice throughout the complete duration of the experiment. Comparison between groups was assessed with an unpaired two-tailed T test. ns: non-significant. <u>Abbreviations:</u> GSK3β… glycogen synthase kinase 3 beta; CLP.. cecal ligation and puncture; EE.. energy expenditure; VO2.. oxygen consumption; RER.. respiratory exchange ratio; m.. meters.

**Figure S13.**
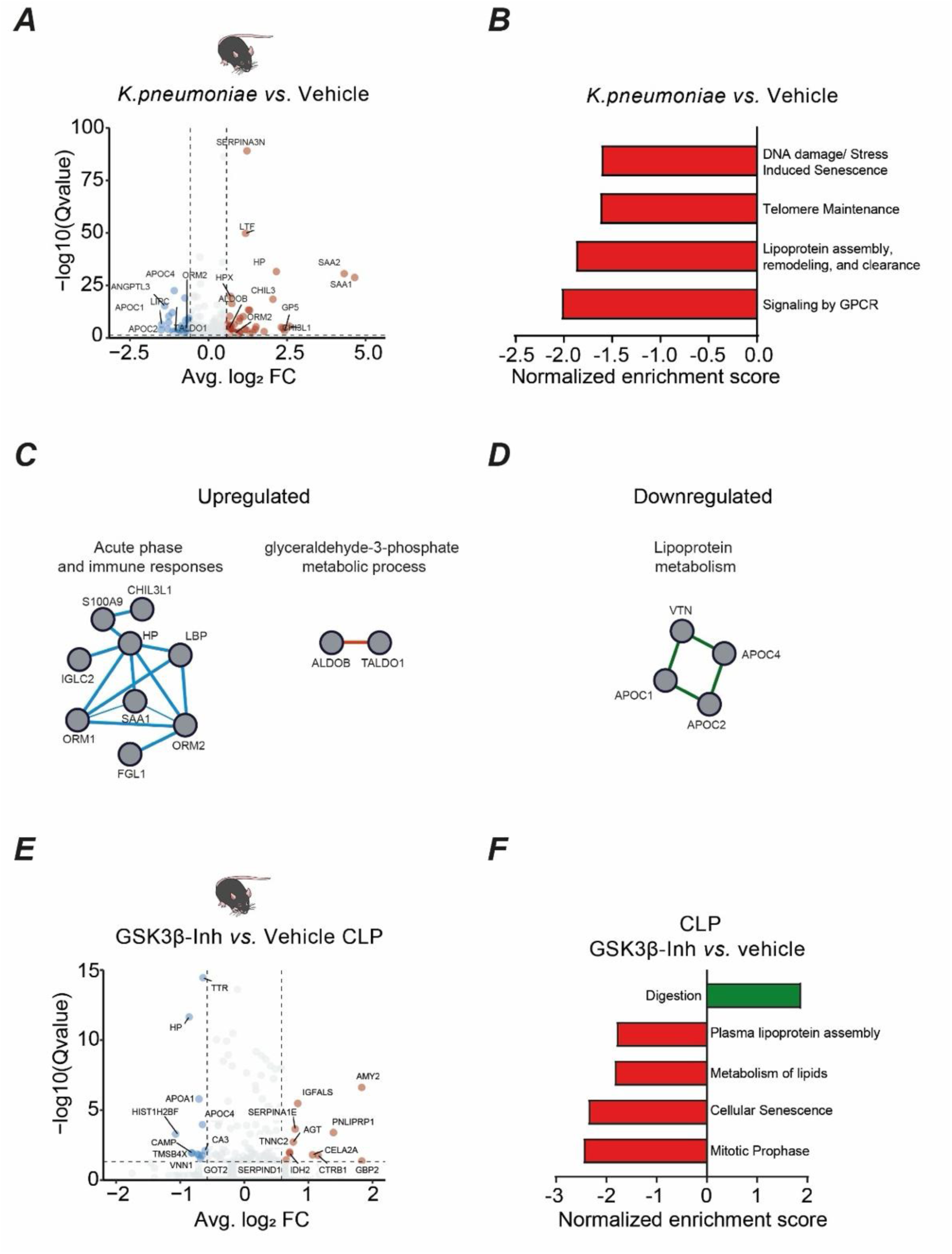
Hypometabolic-associated plasma proteomics correlates to human sepsis severity. Related to Fig. 4. **(A)** Volcano plot depicting differentially expressed proteins in plasma from mice infected with *K.pneumoniae* compared to mice treated with vehicle. **(B)** Gene-set enrichment analysis (GSEA) of differentially expressed proteins in plasma from mice infected with *K.pneumoniae* compared to mice treated with vehicle. Enriched protein-protein interaction modules from differentially **(C)** upregulated and **(D)** downregulated proteins that are shared between *K.pneumoniae* infected mice with human sepsis datasets. **(E)** Volcano plot depicting differentially expressed proteins in plasma from CLP mice subjected to GSK3β compared to CLP mice treated with vehicle. **(F)** GSEA of differentially expressed proteins in plasma from CLP mice subjected to GSK3β compared to CLP mice treated with vehicle.

## Notes

### Competing Interest Statement

The authors have declared no competing interest.

